# A domain-swapped proPC1/3 structure reveals a conformational checkpoint for ER export

**DOI:** 10.64898/2026.07.21.739776

**Authors:** Pankaj Kumar, Phillip A. Hyon, Rishi Nixon, Ashley V. Bullington, Ciara A. Payne, Iris Lindberg, Daniel L. Kober

## Abstract

Prohormone convertase 1/3 (PC1/3, encoded by *PCSK1*) is a serine protease expressed in neuroendocrine cells that is required to produce insulin, glucagon-like peptide-1, adrenocorticotrophic hormone, and other peptide hormones. PC1/3 is synthesized as the zymogen proPC1/3, which undergoes autocatalytic maturation in the endoplasmic reticulum (ER) before trafficking to secretory granules. However, how autocatalytic maturation licenses the ER exit of PC1/3 remains unknown. Here, we determined the structure of an immature, catalytically inactive human proPC1/3^S382A^, which exits the ER as a domain-swapped homodimer. This domain-swapped conformation allows proPC1/3^S382A^ to complete a conserved calcium pocket normally formed after autocatalysis and thereby become competent for ER exit. By defining this calcium pocket as a conformational checkpoint for ER export, our findings provide insights into the maturation of PC1/3 which may apply broadly to the PCSK family. Finally, our structure provides explanations for the functional consequences of many deleterious *PCSK1* mutations.

## Introduction

Plasma glucose levels are regulated in large part by the competing actions of glucagon and insulin, hormones that are produced from longer precursors in specialized secretory cells in the pancreas. While multiple enzymes contribute to hormone maturation, the proteases making the determinative cuts to produce insulin and glucagon belong to the family of proprotein convertase subtilisin/kexin proteases (PCSKs).^1^ The PCSK family contains nine members in humans, seven of which are closely related to the yeast Kex2 and cleave substrates with the general motif of (K/R)_4_-(X)_3_-(K/R)_2_-(R)_1_↓.^2^ The PCSKs play diverse roles across biology, processing many peptide hormone precursors, growth factors, transcription factors, and other substrates. Several PCSK family members are also exploited by viruses, including SARS-CoV-2,^3^ and have been proposed as potential targets for antivirals and cancers.^2^ Despite their importance across human biology, how PCSKs are regulated remains poorly understood.

PCSKs are matured through a conserved mechanism. The peptidase domain folds as a zymogen in the endoplasmic reticulum (ER). Folding is templated by an inhibitory prodomain, and this zymogen is referred to as a proPC (ex: proPC1/3). The peptidase domain folds with the primary cleavage motif from the prodomain trapped in the peptidase active site. This holds the enzyme inactive until it acquires a conformation that is competent to cleave its prodomain through an autocatalytic mechanism. Following cleavage, the cleaved prodomain (also called the propeptide) remains noncovalently associated with the enzyme (the PC). This is the state in which all PCSKs except PC2 are thought to exit the ER^2^. However, the folding and maturation of the PCSK family remains poorly understood. This is due in part to the absence of structures for an immature kexin or PCSK; the closest example is the distantly related pro-Kumamolysin.^4^ Prohormone convertase 1/3 (PC1/3, also known as Neuroendocrine convertase 1, NEC1, encoded by *PCSK1*) is one of two “prohormone convertases” (the other being PC2, encoded by *PCSK2*), because it is predominantly expressed in endocrine, neuronal, and neuroendocrine tissues. PC1/3 accumulates in regulated acidic secretory granules where it processes critical substrates such as proopiomelanocortin (POMC) in the production of adrenocorticotrophic hormone (ACTH), proglucagon in the production of glucagon-like peptide-1 (GLP-1) and GLP-2, and proinsulin in the production of insulin, amongst others.^5, 6^ While PC1/3 and PC2 share some substrates, they have critical non-redundant roles owing to distinct expression patterns^7, 8^ and distinct substrate preferences.^9, 10^ In animals, PC1/3 is critical for insulin production in β cells, whereas PC2-deficient animals lack glucagon but show minimal loss of insulin.^5, 6^

PC1/3 has strong genetic links to disease. A variety of *PCSK1* mutations are associated with diabetes and obesity, among other pleiotropic effects.^7^ The *PCSK1* G593R variant was among the first examples of a point mutation linked to obesity in humans.^11^ In infants, inactivating mutations cause severe intestinal malabsorption, diarrhea, and can be fatal.^11^ PC1/3 variants are found in up to 25% of the population, and represent the third most common monogenic contributor to obesity across various human populations.^11^ Certain PC1/3 mutations are known to impair protein stability and trafficking, while others are secreted at normal levels and have only modest effects on activity.^12^ PC1/3 can oligomerize in cells,^10^ and mutations that impair PC1/3 trafficking can have a dominant effect on wild-type (WT) PC1/3.^13^ However, except for the inactivating mutations, the structural basis for the biological effects of these mutants is not clear. Despite intensive crystallographic efforts by several laboratories, structural insights into the prohormone convertases are lacking.^14^

Here we present the structure of human proPC1/3, trapped using a catalytic mutation (S382A). ProPC1/3^S382A^ forms an unusual domain-swapped dimer, where the prodomain of one molecule inhibits the peptidase domain of the other. Interestingly, this conformation specifically licenses its ER exit, and our data reveal that the gatekeeping step for the ER exit of PC1/3 is not autocatalysis, but the completion of a conserved calcium binding site. In furin, and potentially in other PCSKs, the formation of this site requires autocatalysis to liberate an Asp residue to chelate the calcium ion. By folding as a domain-swapped homodimer, the inert proPC1/3^S382A^ acquires this key feature of a mature PC1/3. Additionally, our structure of proPC1/3^S382A^ explains the deleterious effects of many loss-of-function disease variants. Our findings provide insights into the folding and maturation of PC1/3, which may apply to the broader PCSK family.

## Results

### A dimeric proPC1/3 can exit the ER

In order to define the general properties required for maturation of a proPC into a PC, we decided to revisit the question of whether proPCs are generally prohibited from exiting the ER. ER exit is generally considered to be the rate-limiting step for the constitutive secretion of most proteins,^15^ and HEK 293 cells possess only a constitutive secretory system, i.e. lack regulated secretory granules.^16^ Therefore, HEK 293T cells offered a facile means to test for ER exit of proPCs. We tested FLAG-tagged wild-type (WT) and catalytic mutant versions of four kexin-like PCSKs: human PC1/3, human furin, mouse PC5A, and human PC7, spanning the diversity of this family (**Fig. 1a** and **Supplemental Fig. 1a**). PC2 was not tested because of its known dependence on its chaperone 7B2 for ER exit, suggesting a unique ER exit mechanism.^17, 18^ For this assay, furin and PC7 were subcloned without their C-terminal transmembrane helices to enable secretion. HEK 293T cells were transfected with these plasmids, after which the cell lysates and conditioned media were probed by immunoblot (**Fig. 1b**). Both the mature PC1/3 and catalytically inert proPC1/3^S382A^ were readily detected in the medium, indicating that both can exit the ER. In contrast, the corresponding mutants of mouse PC5A (S388A), furin (D153A), and PC7 (D187A) were not secreted (**Fig. 1b**). HEK 293T cells also retained the potentially misfolded PC1/3 G593R mutant, consistent with our previous report^13^ (**Supplemental Fig. 1b**). In order to confirm these results in a neuroendocrine cell line with a regulated secretory pathway, we tested the ability of these same PC1/3 forms to traffic in AtT-20 cells, a model for the study of peptide hormone maturation for the past four decades.^19^ Immunofluorescence-based imaging of AtT-20 cells transfected with FLAG-tagged WT PC1/3 or proPC1/3^S382A^ revealed that both species can co-localize with markers for the TGN (**Fig. 1c**) and also appeared at the tips of the cells, where secretory granules are found. ^20^ These data show that proPC1/3^S382A^ and WT proPC1/3 can adopt a folded state that is licensed for ER exit.

**Figure 1:**
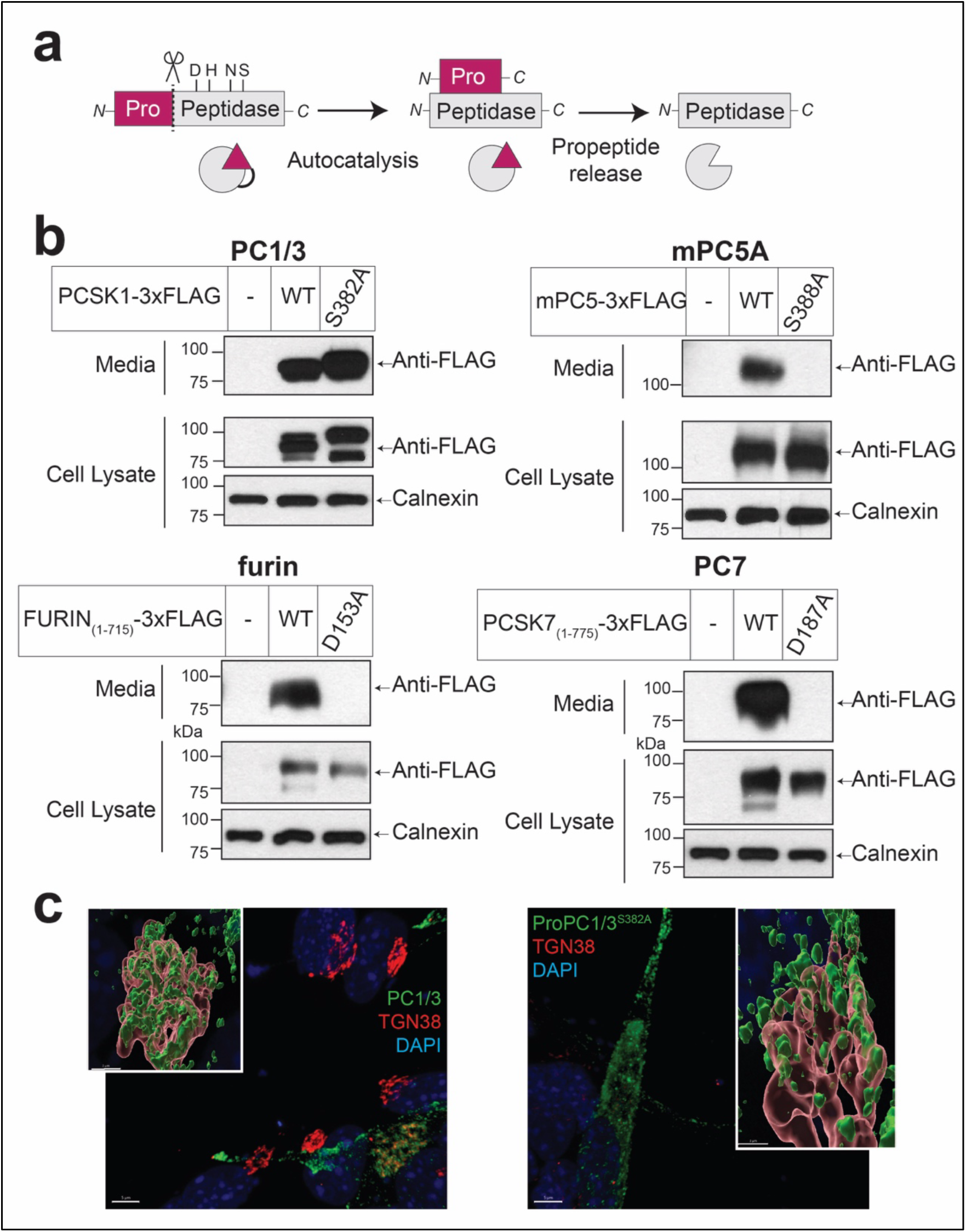
ProPC1/3^S382A^ can exit the ER. **a)** Schematic for PCSK maturation and activation. Following cleavage of the signal peptide, PCSKs have an N-terminal prodomain that templates the folding of the peptidase domain. The peptidase domain houses the Asp, His, and Ser catalytic triad, and an oxyanion hole residue, typically an Asn. Following autocatalytic maturation, the cleaved propeptide must be released. **b)** Expression and secretion of various PCSKs. HEK 293T cells were set up in 12-well dishes at a density of 400,000 per well. The next day, cells were transfected and processed as described in Methods. Each well was transfected with 1 µg of empty pEZT vector (lane 1) or the indicated plasmid (lanes 2-3). Samples were electrophoresed on 10% SDS-PAGE and subjected to immunoblot analysis for anti-FLAG or anti-Calnexin. **c)** Staining of transfected PC1/3 and proPC1/3^S382A^ in AtT-20 cells. AtT-20 cells were transfected with PC1/3-3xFLAG or proPC1/3^S382A^-3xFLAG, fixed with paraformaldehyde after 24 h, and immunostained for FLAG (green) and TGN38 (red), z-stacked on a confocal microscope, and segmented for rendering in 3D using Imaris. Scale bars are 5 µm (2D scan) and 3 µm (inset).

ProPC1/3 experiences several cleavage events as it traffics through a regulated secretory pathway.^11^ Autocatalytic maturation in the ER by cleavage at the primary site produces the so-called “87 kDa” form of PC1/3 (**Fig. 2a**).^21^ Like furin, the 87 kDa PC1/3 releases its cleaved prodomain after a secondary cleavage of the prodomain in the TGN.^22, 23^ In cells with a regulated secretory pathway, PC1/3 undergoes additional processing steps in the acidic secretory granules, where it intermolecularly cleaves off its C-terminal domain to produce the “66 kDa” form of PC1/3.^24^ The 66 kDa form is the major species of PC1/3 in neuroendocrine cells.^25^ To purify intact 87 kDa PC1/3 or proPC1/3^S382A^, we utilized C-terminal FLAG tags (**Fig. 2a**) and purified these proteins from the conditioned medium of BacMam-infected GnTI-deficient HEK 293S cells^26^ (**Fig. 2b**). Following anti-FLAG affinity chromatography, PC1/3 eluted from gel filtration in two peaks, which we termed Peak I and Peak II. Coomassie staining revealed that Peak I contained PC1/3 in a non-covalent heterodimer with its cleaved propeptide, whereas Peak II contained 87 kDa PC1/3 without the propeptide (**Fig. 2c**). Analysis of an independent purification showed that Peak II contained a small amount of the apparent 66 kDa species and a FLAG-reactive band of approximately 21 kDa, consistent with the cleaved C-terminal domain (**Supplementary Fig. 2a**).

**Figure 2:**
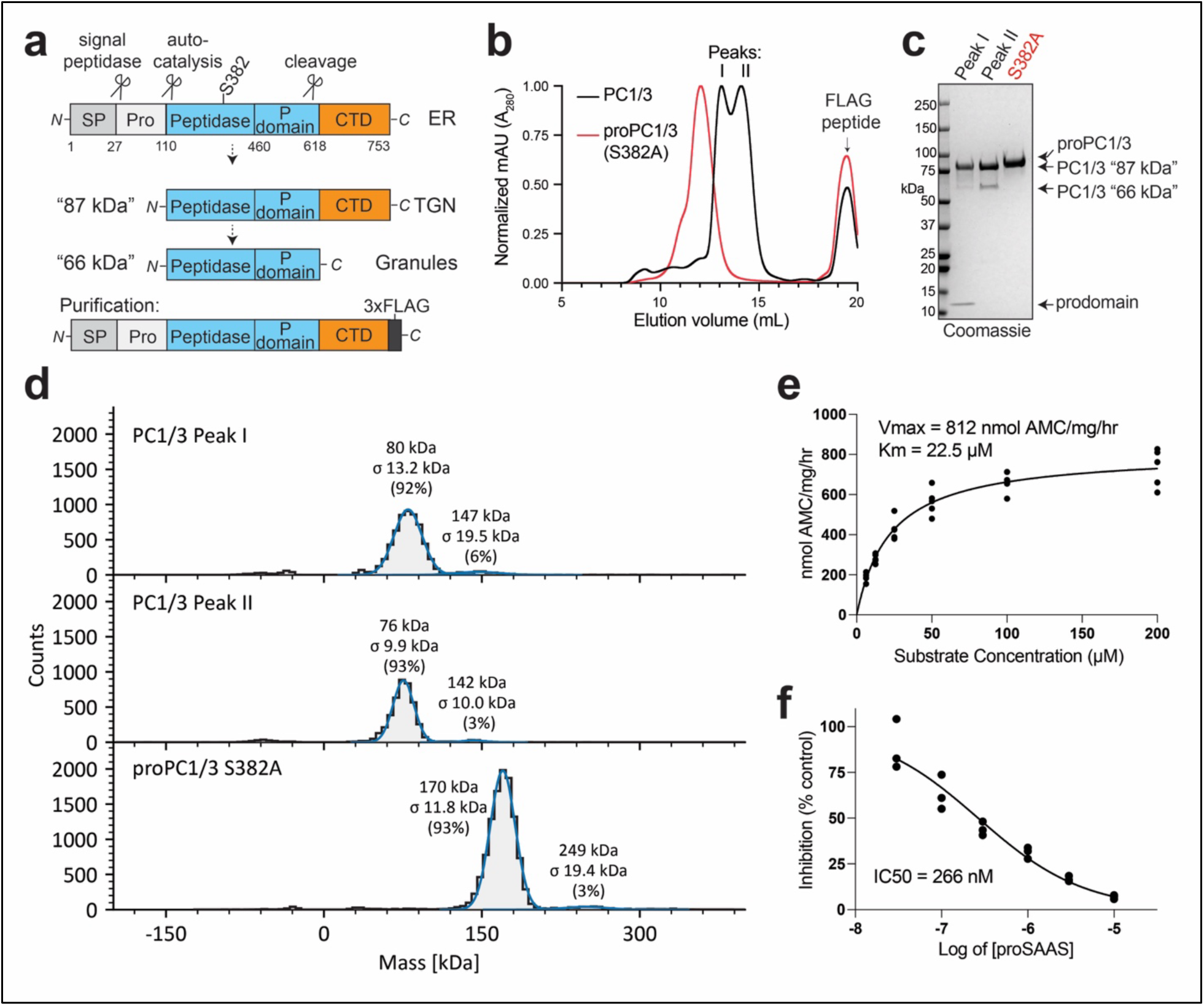
Characterization of PC1/3 and proPC1/3^S382A^. **a)** Schematic for the stages of PC1/3 maturation and the purification construct. **b)** Normalized gel filtration chromatograms for the purification of proPC1/3^S382A^ (red trace) and PC1/3 (black trace). The anti-FLAG eluates were concentrated and subjected to gel filtration over an equilibrated Superdex 200 Increase 10/300 column at 4°C a flow rate of 0.7 mL per min as described in Methods. The void fraction on this column is ∼8.5 mL. **c)** Coomassie-stained 4-15% SDS-PAGE gel with 1.7 µg of sample from the traces in B. **d)** Mass photometry measurements of the samples from B. **e)** Dose-response fluorogenic protease assays. Assay contained 50 nM enzyme incubated with the indicated concentration of pyr-RTKR-AMC, as described in Methods. Each point is the mean of three technical replicates. The assay was conducted five times using protein from two independent purifications (two replicates from one purification and three replicates from the other). Data were fit using Michaelis-Menten analysis with GraphPad Prism. **f)** Inhibition of 50 nM PC1/3 by mouse proSAAS CT peptide using the fluorogenic peptide assay as described in Methods. Each point is the mean of three technical replicates (n=3). Inhibition curve fit done with GraphPad Prism. Pro = prodomain; CTD = C-terminal domain; ER = endoplasmic reticulum; TGN = Trans Golgi Network.

In contrast to PC1/3, the inert mutant proPC1/3^S382A^ eluted earlier on gel filtration and contained a single major peak (**Fig. 2b**). This proPC1/3^S382A^ did not contain free prodomain or a 66 kDa species (**Fig. 2c** and **Supplementary Fig. 2a**). Mass photometry measurements showed that the proPC1/3^S382A^ formed a dimer (experimental mass: 170 ± 11.8 kDa, theoretical monomer = 84.4 kDa) whereas PC1/3 had masses of 80 ± 13.2 kDa and 76 ± 9.9 kDa, within expectations for PC1/3 ± its cleaved propeptide (theoretical values PC1/3 = 74.6 and propeptide = 9.8 kDa) (**Fig. 2d**). These solution measurements demonstrate that proPC1/3^S382A^ is secreted in a different conformation than WT PC1/3.

Next, we tested whether our FLAG-tagged PC1/3 retained its expected enzymatic activities. First, PC1/3 was able to mature into its 66 kDa form in a pH-dependent manner (**Supplementary Fig. 2b**). As expected, PC1/3 also processed proglucagon more efficiently at pH 5.5 than pH 7.0 whereas proPC1/3^S382A^ had no activity (**Supplementary Fig. 2c**). (Proglucagon contains multiple dibasic motifs and two confirmed PC1/3 cleavage sites,^27, 28^ but we did not identify the product peptides in this assay). In these proglucagon cleavage assays, we observed some product formed at pH 7 by the Peak II sample, so we then carried out pH-dependence studies with the Peak I and Peak II PC1/3 using the fluorogenic reporter substrate, pyroglutamyl (pyr)-Arg-Thr-Lys-Arg-AMC (pyr-RTKR-AMC). With this substrate peptide, we measured similar activities for each peak at pH 5.5 and a bimodal activity range for Peak II, which had a second activity maximum around pH 7 (**Supplementary Fig. 2d**). Based on these results, we combined Peak I and Peak II and carried out a Michaelis-Menten substrate concentration assay at pH 5.5, corresponding to acidic secretory granules. In these conditions, PC1/3 had an apparent K_M_ of ∼22 µM for Pyr-RTKR-AMC and a Vmax of ∼800 nmol product per milligram per hour, in general agreement with previous reports of 134-480 nmol product per milligram per hour,^10^ (**Fig. 2e**). Importantly, the activity of 50 nM PC1/3 could be inhibited by pre-incubating the PC1/3 with a peptide containing the residues 219-258 of mouse proSAAS,^29^ a known potent inhibitor of PC1/3^29–31^ (**Fig. 2f**). Collectively, these results validate our epitope-based purification as a facile means to enrich well-behaved, full-length PC1/3 and dimeric proPC1/3^S382A^.

### The proPC1/3^S382A^ forms a domain-swapped homodimer

We used cryogenic electron microscopy (cryo-EM) to determine the structure of the proPC1/3^S382A^ dimer to a nominal resolution of 2.9Å (**Supplementary Fig. 3** and **Supplementary Table 1**). Surprisingly, proPC1/3^S382A^ forms a domain-swapped homodimer where the prodomain of one molecule is bound to the peptidase domain of the other molecule (**Fig. 3a**). There is unambiguous density from the prodomain up to R114, flexible but traceable density for residues 115-119, with strong density resuming at L120. Density for an N-linked glycan attached to N173 further supports the register assignments (**Fig. 3b**). The map quality supported modeling three calcium ions and one sodium ion that bind in conserved pockets. Lower local resolution for the prodomain precluded modeling a reported sodium site.^32^ We could not resolve any of the C-terminal domain, suggesting flexibility. Interestingly, the proPC1/3 dimer interface uses a surface that is structurally analogous to the one used by PCSK9 to bind the EGF-A domain of the LDL receptor, potentially highlighting a broad role for this surface in protein-protein interactions among the PCSKs (**Supplementary Fig. 4a**).

**Figure 3:**
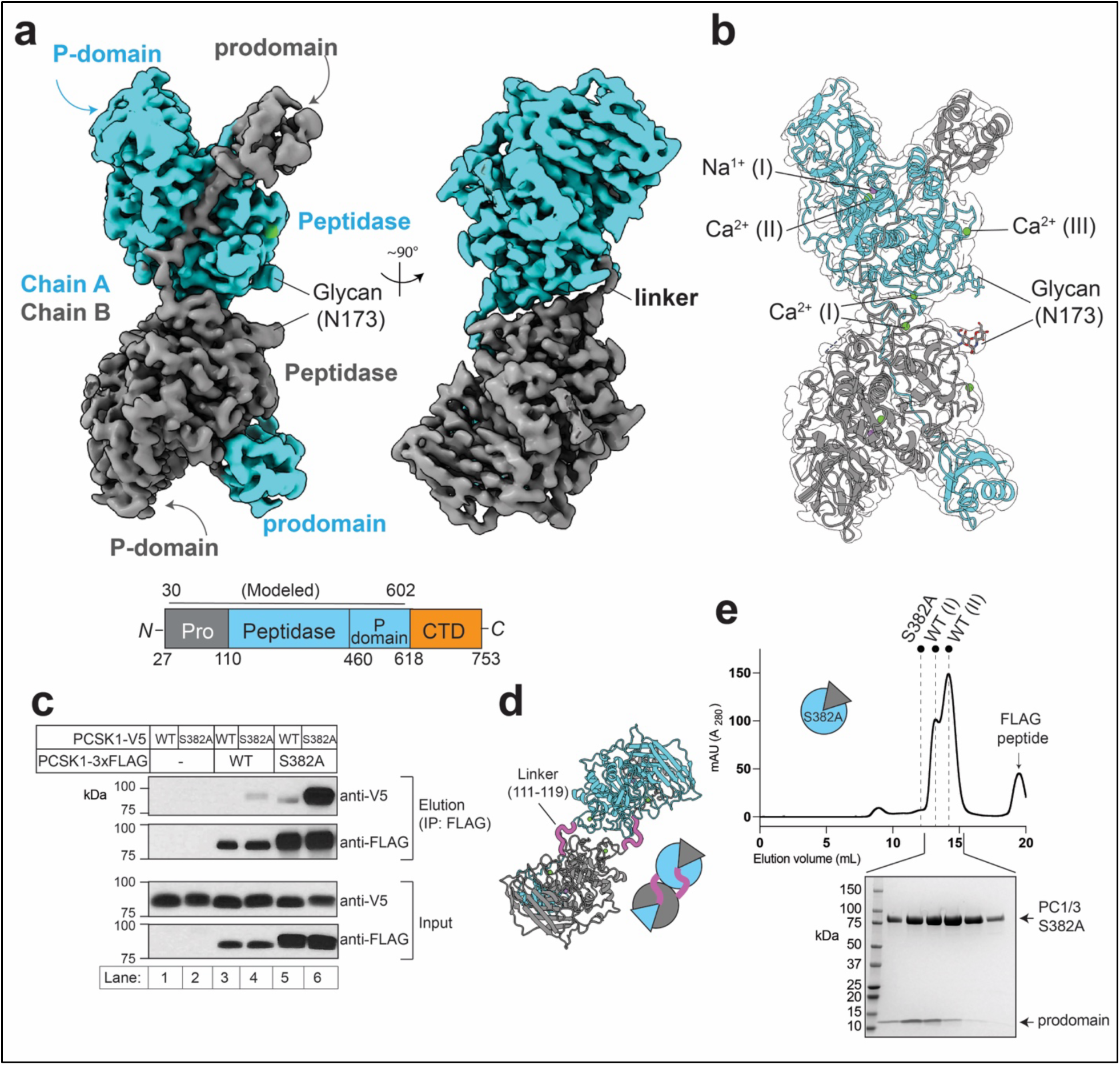
Cryo-EM structure and functional analysis of the proPC1/3^S382A^ dimer. **a)** Cryo-EM map of the proPC1/3^S382A^ dimer in two orientations and colored by protein chain. Unsharpened map shown at contour 0.00487. Domains of proPC1/3 are labeled. The schematic of proPC1/3 domains depicts the region resolved in the single-particle reconstruction. **b)** Atomic model for proPC1/3^S382A^ shown in transparent, unsharpened map. The glycan tree attached to N173 is labeled, as are the modeled calcium sites I-III and sodium ions. **c)** Co-immunoprecipitation assays testing for an interaction between orthogonally tagged PC1/3 proPC1/3^S382A^. HEK 293T cells were set up and transfected with 3 µg each of the indicated PCSK1 plasmids. Empty pEZT vector was used to adjust the total DNA amount to 6 µg. Following expression for two days, conditioned supernatants were subjected to pulldown with anti-FLAG resin, as described in Methods. Samples were electrophoresed on 10% SDS-PAGE and immunoblotted for the indicated epitopes. **d)** Model of dimeric proPC1/3^S382A^, with the linker residues colored as magenta tubes. **e)** Expression of PC1/3^ecto^ (S382A) in trans with the PC1/3 prodomain produces a PC1/3^S382A^ that behaves like WT PC1/3 on gel filtration. Gel filtration trace for the PC1/3^ecto^ (S382A)-prodomain complex, with markers indicating the elution volumes of the full-length proPC1/3^S382A^ and PC1/3 (from Fig. 2). Inset is Coomassie-stained 4-15% SDS-PAGE gel showing the fractions highlighted with brackets. Panels A, B, and D were generated using ChimeraX.

The proPC1/3^S382A^ prodomain, peptidase domain, and P-domain, all adopted their typical folds; therefore, we focused on validating the dimer interaction. We conducted co-immunoprecipitation (co-IP) experiments in which FLAG-tagged PC1/3 or proPC1/3^S382A^ were used as bait for V5-tagged PC1/3 or proPC1/3^S382A^. While a weak interaction was observed between PC1/3 and proPC1/3^S382A^ (**Fig. 3c**, lanes 4-5), the strongest interaction was observed when both constructs harbored the S382A mutation (**Fig. 3c**, lane 6). Additionally, the proPC1/3^S382A^ co-IP results were similar when the proPC1/3s were co-expressed with His-tagged human proSAAS (**Supplementary Fig. 4b**) suggesting that proSAAS does not play a role in dimer formation.

The proPC1/3^S382A^ dimer interface is large and encompasses 3610 Å^2^ of buried surface area, according to analysis on the PISA server.^33^ Most of the interaction is between the prodomain and peptidase domains. The interface between the peptidases contains only ∼420 Å^2^. This analysis suggested that the linker region between the prodomain and peptidase is critical for the stability of the proPC1/3^S382A^ dimer (**Fig. 3d**). To test this, we generated a pair of constructs encoding either the prodomain alone or the FLAG-tagged ectodomain of PC1/3 without its prodomain, hereafter PC1/3^ecto^ (**Supplementary Fig. 4c**). Secretion of PC1/3^ecto^ (WT) or PC1/3^ecto^ (S382A) could be rescued by co-expressing the prodomain *in trans* (**Supplementary Fig. 4d**), similar to previous studies with PCSK9.^34^ To test whether PC1/3^ecto^ (S382A) could form a stable dimer with a second PC1/3^ecto^ (S382A), we co-expressed PC1/3^ecto^ (S382A) with its untagged prodomain, purified the secreted complex using anti-FLAG affinity resin, and found that it behaved similarly to WT PC1/3 on gel filtration (**Fig. 3e**). Mass photometry recorded masses of 85 kDa and 79 kDa for material collected from the two shoulders, indicating that PC1/3^ecto^ (S382A) does not form stable dimers (**Supplementary Fig. 4e**). These data suggest that the linker region is responsible for maintaining the dimeric complex.

Our structure of a proPC1/3 ectodomain with a domain-swapped prodomain closely resembles a recent crystal structure of matured furin in complex with a PC1/3 prodomain (PDB 9FIE,^32^ **Fig. 4a**, RMSD = 0.785 Å). Further, we observed only minor differences between our prodomain and a crystallized mutant prodomain of PC1/3 (PDB 9FIC,^32^ **Supplementary Fig. 5**). Our structure shows proPC1/3 bound to its primary substrate motif and captures interactions before and after the cleavage site (**Fig. 4b**). The canonical primary cleavage motif R_107_-S_108_-K_109_-R_110_ ↓ is well-resolved and both the catalytic residues (sans the nucleophilic side chain of S382) and the oxyanion hole residue are in their expected conformations (**Fig. 4c**). The basic P4, P2, and P1 residues all fit into acidic pockets, with each making hydrogen bonds with the peptidase (**Fig. 4d-f**). The S1 pocket is the deepest, provides the most polar contacts, and is stabilized by the calcium site-(II) ion. The S2, and S4 pockets are each buttressed by W268, which is analogous to W254 of furin,^35^ W273 of kexin, and I308 of S1P. This conserved hydrophobic residue controls the accessibility of the substrate pockets of S1P^36^ and can be displaced by furin inhibitors that bind through an induced-fit mechanism.^37^

**Figure 4:**
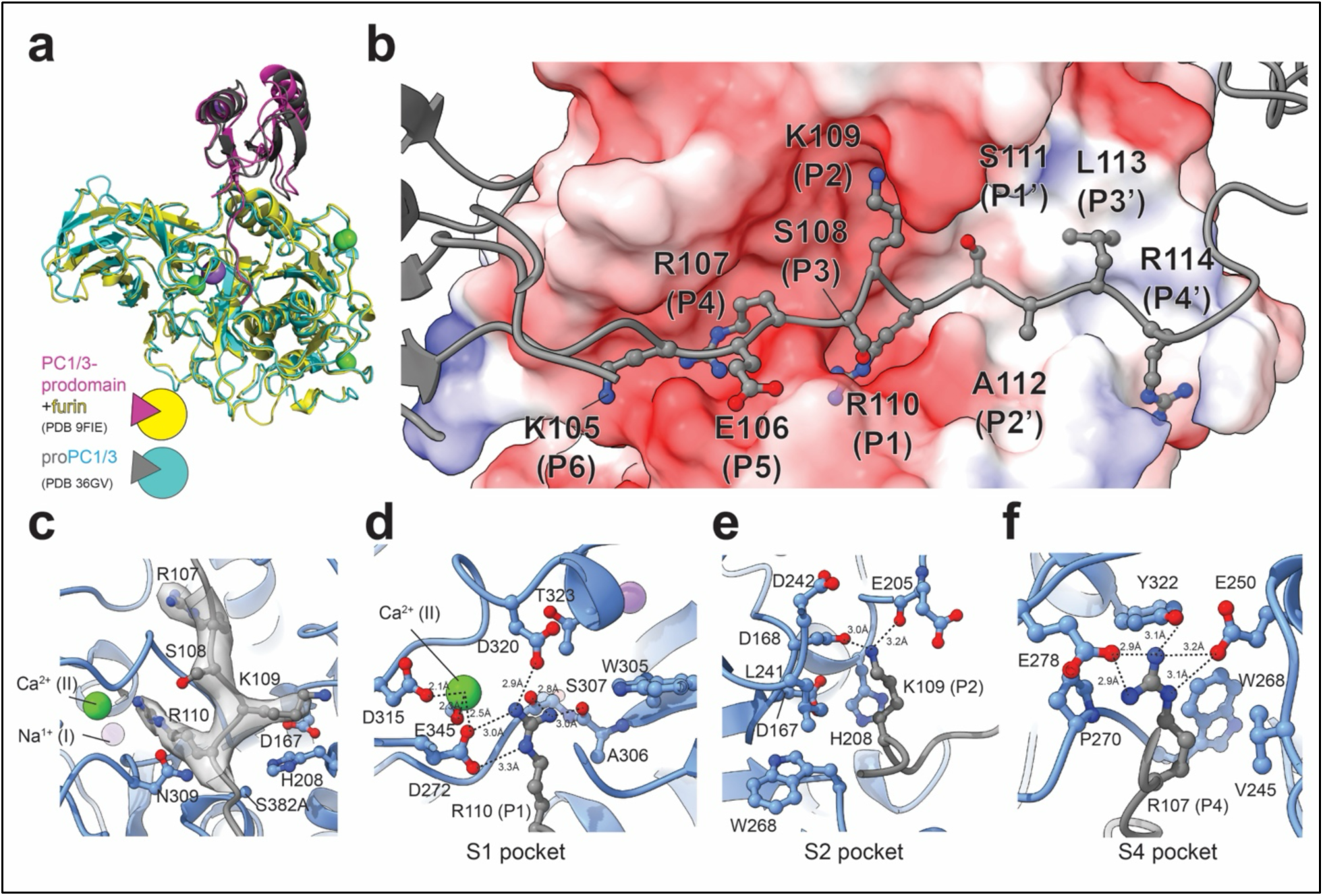
The proPC1/3 prodomain as a model for substrate recognition by PC1/3. **a)** Superposed structures of one copy of the proPC1/3^S382A^ complex with the furin-PC1/3 prodomain complex, PDB 9FIE. Secondary structure cartoons are colored as indicated in schematic. **b)** The prodomain cleavage motif bound to proPC1/3^S382A^. Prodomain is shown as grey cartoon, with residues 105-114 shown as sticks and colored by heteroatom. Peptidase domain surface is colored by electrostatics, with red =-10 and blue = +10 kT/e. **c)** Primary cleavage motif and catalytic residues. Prodomain shown as grey cartoon with residues 107-110 shown as sticks and colored by heteroatom. Peptidase domain shown as blue cartoon. Catalytic residues and oxyanion hole residues shown as sticks and colored by heteroatom. Sharpened map is shown as transparent grey surface contoured at level 0.0139 and carved at 2Å around residues 107-110. **d-f)** Structural models showing interactions at the S1, S2, and S4 pockets and depicted as in panel C. Potential polar contacts are highlighted with dashed lines. Panel A was generated using Pymol; the others were generated using ChimeraX.

PC1/3 and PC2 differ in substrate preferences at the P1’ and P2’ positions (P’ denotes residues C-terminal to the cleavage site), with PC1/3 unable to cleave substrates with large aliphatic residues at these positions, unlike PC2,^7^ a trend confirmed in studies of animal peptide hormones^38^. Our structure suggests why PC1/3 disfavors certain P’ residues. Following the scissile bond, P1’ residue S111 makes a hydrogen bond with the catalytic residue H208 and P4’ residue R114 makes a salt bridge with E359. R114 may also make cation-pi stacking interactions with W342 and Y360. In our structure, the S1’ pocket is not large and would not accommodate a bulky residue. The S2’ pocket is less restrictive but is more negatively charged, likely disfavoring the aromatic and hydrophobic residues accommodated by PC2.

At least 35 coding and noncoding variants have been identified in *PCSK1* ^39^, many of which are associated with obesity or related diseases. Our experimental structure reveals the location of several residues implicated in diseases and provides explanations for their biological effects. For this analysis, we focused on the variants demonstrated to produce deleterious biochemical effects (**Fig. 5a, b**).^7, 11, 39^ Three Gly to Arg substitutions have been shown to prevent autocatalytic maturation and increase ER retention. G593R, the first identified, was found in a patient with childhood obesity and disrupted endocrine functions.^40^ The G593R variant, as well as a G209R variant found in a patient with a similar phenotype,^41^ can interact with WT PC1/3 and reduce the secretion and activity of WT PC1/3 when co-expressed.^13^ G593 is located in the P-domain, at the end of one of the major beta strands. An Arg mutation at this position would project into the P domain and disrupt the hydrophobic packing (**Fig. 5c**). The G209R mutation would disrupt the positioning of the catalytic D167, precluding catalysis (**Fig. 5d**). The third pathogenic Gly to Arg mutation at position 226 was not secretable.^42^ This residue lies close to the calcium-I site and the intrusion of an Arg would disrupt folding, potentially preventing a disulfide bond between C225 and C374 (**Fig. 5e**). Additionally, a recent report identified a V229F variant in an adolescent patient with obesity and type-2 diabetes.^43^ The V229 residue is buried within the core of the peptidase domain, and a Phe substitution is unlikely to be tolerated due to steric clashes (**Fig. 5f**).

**Figure 5:**
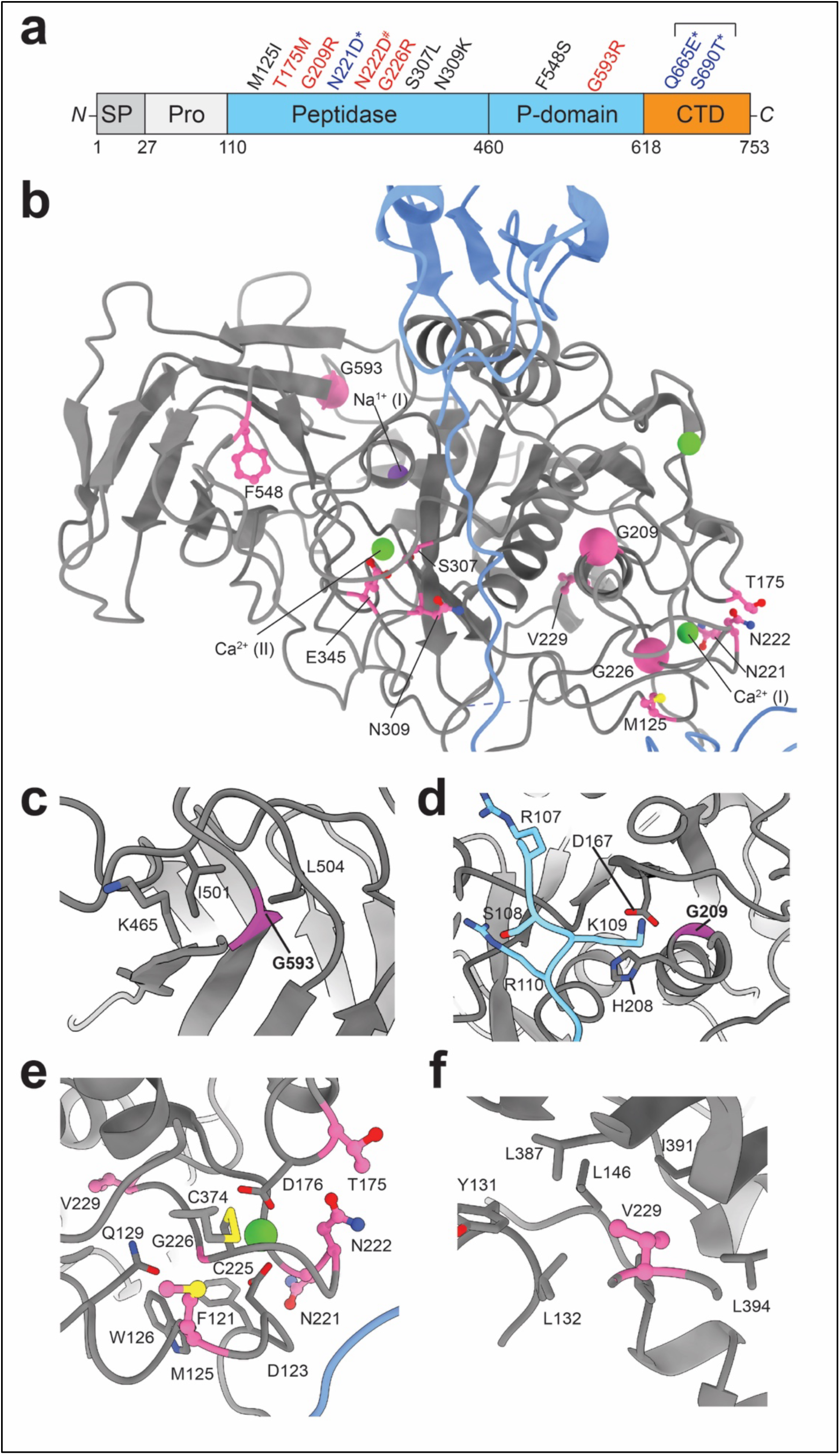
Structural basis for loss of function PC1/3 disease variants. **a)** Locations of disease-associated variants in PC1/3. Schematic depicts the domain boundaries of PC1/3 with approximate location of disease variants. Variants in black text have been shown to allow autocatalysis and ER exit but reduced activity. Variants in blue are common polymorphs. Variants in red reduce autocatalysis and/or prevent ER exit. # indicates the loss of function N222D variant produced in mice. The bracket highlights Q665E and S690T, which are in linkage disequilibrium. **b)** Position of disease-associated residues on proPC1/3^S382A^. Key side chains shown in sticks. C-alpha atoms of glycine residues are shown as spheres. **c-f)** Local environment for G593 (c). G209 (d), G226 and calcium site-I (e), and V229 (f). Panels b-f generated with ChimeraX. SP = signal peptide, Pro = prodomain, CTD = C-terminal domain.

Additional disease-associated variants, (M125I, T175M, and N221D) cluster around the calcium-I site, as previously suggested from homology modeling (**Fig. 5e**).^7, 42^ The T175M variant also disrupts the glycosylation motif for N173, and this mutation reduces the maturation and secretion of PC1/3.^42^ The M125I variant was reported to reduce enzymatic activity by ∼75% but did not prevent autocatalytic maturation or secretion. This residue is near the calcium site and is part of the dimer interface,^42^ highlighting the importance of these regions in PC1/3’s enzymatic activity (**Fig. 5e**).

The three most common PC1/3 variants are N221D, Q665E, and S690T. ^7^ The latter two residues, which were not resolved in our structure, are in linkage disequilibrium and are strongly associated with human obesity, but did not show obvious deficiencies in vitro.^13^ The N221D variant was reported to have a modest ∼30% reduction in activity,^13^ but was less stable.^44^ Additionally, forward genetics in mice produced a N222D mutation that causes obesity^45^. In vitro, this mutation exhibited greatly reduced autocatalytic maturation, secretion, and increased proteasomal degradation.^12, 46^ Although their side chains do not appear to play direct roles in calcium chelation, the N221D and N222D mutations may alter the region around the calcium site-I (**Fig. 5e**). Finally, an E345A variant was identified in an infant with malabsorptive diarrhea and produced a protein that could undergo autocatalysis and secretion but was inactive against intermolecular substrates. The E345A mutation removes a side chain involved in chelating the calcium site-II ion that supports the S1 pocket, explaining its deficient activity (**Figs. 4d** and **5b**).

### A conserved calcium site licenses the ER exit of proPC1/3

The original crystal structure of furin (PDB 1P8J)^47^ captured the enzyme in its mature state, with a highly conserved calcium binding site-I completed by an Asp that follows shortly after the prodomain at position 115 (**Supplementary Fig. 6a**). It was appreciated that autocatalysis must first occur for D115 to reach this calcium site, since it could not reach this position if the prodomain were intact, i.e., with the primary cleavage motif within the active site.^14^ Because there are no structures of the furin zymogen, we used the AlphaFold 3 server to model furin with the prodomain bound in a pre-catalytic conformation (**Fig. 6a**, *left panels*). Crystal structures such as 9FIE^32^ and the high-resolution 5JXG^35^ captured mature furin with and without a cleaved propeptide, respectively (for 9FIE, the prodomain of PC1/3 was added following purification^32^). In each case, furin D115 chelates calcium site-I, a rearrangement that moves the D115 ∼20Å from its predicted initial position (**Fig. 6a**, *middle and right panels*). Interestingly, in our structure of proPC1/3^S382A^, the domain-swapped architecture enables the homologous residue D123 to chelate the calcium site-I (**Fig. 6b**). With D123 in place, the prodomain extends to interact with the other molecule’s peptidase domain instead of its own. The surprising result is that the proPC1/3^S382A^ mutant is able to chelate the site-I calcium in a manner similar to mature furin (**Supplementary Fig. 6b**).

**Figure 6:**
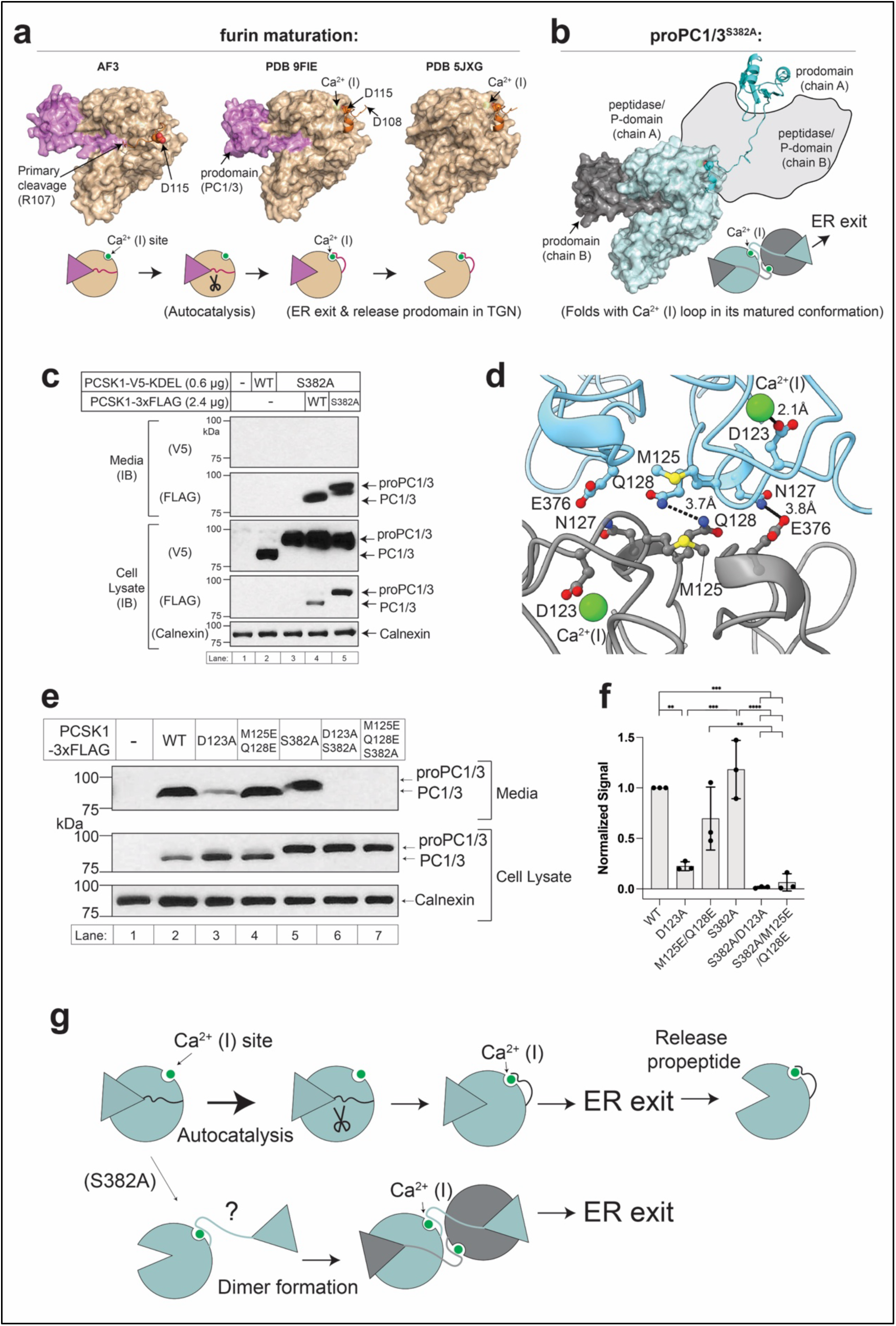
Proposed structural basis for ER exit of proPC1/3^S2382A^: completion of calcium binding site-I. **a)** The different stages of furin maturation. The furin or PC1/3 prodomains are shown in magenta with the linker shown in brown. From left to right, the structural models of furin are derived from AlphaFold server (AF3), PDB 9FIE, and PDB 5JXG. **b)** The proPC1/3^S382A^ dimer enables the completion of the calcium site-I. For this illustration, the peptidase and P-domains of chain B were replaced with the grey outline. The prodomain of chain B are depicted with grey surface representation and the peptidase and P-domains of chain A are depicted with cyan surface representation. The prodomain and linker of chain A are depicted in cyan cartoon format. **c)** ProPC1/3 is not cleaved *in trans* by PC1/3 in the ER. HEK 293T cells were set up in 6-well plates and transfected with 3 µg of DNA. After expressing for one day, cell and supernatant samples were harvested and processed as described in Methods. IB = immunoblot for the epitope indicated in parentheses. **d)** Structure of the proPC1/3^S382A^ dimer interface, highlighting interactions targeted by mutagenesis. Key residues are shown as sticks and colored by heteroatom. Dashed lines and distances indicate potential polar interactions. **e)** Expression and secretion of PC1/3 and proPC1/3 mutants. HEK 293T cells were set up in 12-well plates and transfected with 1 µg of DNA. Samples were processed and subjected to analysis by immunoblot as described in Methods. ProPC1/3 and PC1/3 were detected with anti-FLAG antibodies. **f)** Quantification of three replicates of the experiment shown in (e). The immunoblot signals for secreted FLAG-tagged proteins were normalized to the signal from WT PC1/3 from each experiment (lane 2). Data were analyzed by one-way ANOVA with Tukey’s correction for multiple comparisons and comparisons with a corrected p value < 0.01 are labeled (** < 0.01, *** < 0.001, and **** < 0.0001). Error bars are SD. **g)** Model for WT proPC1/3 maturation and the ER exit of proPC1/3^S382A^. Panels a and b were generated with Pymol. Panel d was generated with ChimeraX.

Our results suggest two interesting hypotheses. First, that WT proPC1/3 might also mature *in trans* via the domain-swapped configuration; and second, that the completion of calcium site-I enables the ER exit of proPC1/3^S382A^. To test the first hypothesis, we co-transfected the proPC1/3^S382A^-FLAG plasmid with V5-tagged WT PC1/3 or proPC1/3^S382A^ plasmids at a ratio of 1:4, FLAG:V5. However, we only observed a faint band at the expected size for mature PC1/3-FLAG when it was co-expressed with WT PC1/3-V5 (**Supplementary Fig. 6c**, compare lanes 4 and 5). As noted previously (**Fig. 2a**), PC1/3 experiences secondary cleavage of its prodomain *in trans* while traversing the TGN,^22, 32^ a potential complication in this assay (note the doublet form of secreted proPC1/3^S382A^-FLAG in **Supplementary Fig. 6c**). To circumvent the possibility of cleavage in a post-ER compartment, we introduced an ER retention motif (KDEL) onto the C-terminus of V5-tagged proPC1/3 and carried out a similar experiment where V5-tagged proPC1/3^S382A^-KDEL was co-expressed with FLAG-tagged PC1/3 or proPC1/3^S382A^ (**Fig. 5c**). In this format, we also did not observe any evidence for cleavage of proPC1/3^S382A^-FLAG *in trans* by WT PC1/3-V5. Thus, the formation and *in trans* cleavage of a wild-type zymogen dimer is unlikely.

We then tested the second hypothesis, that the formation of the proPC1/3^S382A^ dimer results in the completion of calcium site-I, thus enabling efficient ER exit. Despite some favorable electrostatic interactions, we did not observe hydrogen bond interactions or tight Van der Waals packing in the dimer interface (**Fig. 6d**). Therefore, we targeted this interface through like-charge repulsion by substituting M125 and Q128 for glutamic acid. To test the possible requirement for chelation of calcium site-I by D123, we separately substituted alanine at that position. These mutations were introduced into both the WT and S382A constructs. In HEK 293T cells, neither D123A nor M125E/Q128E prevented autocatalysis when introduced into the WT PC1/3 background, suggesting they do not prevent folding of the peptidase domain (**Fig. 6e**, lanes 1-4). However, in the WT PC1/3 background, the D123A mutation resulted in markedly reduced secretion, whereas M125E/Q128E produced only a modest and non-significant decrease (**Fig. 6e, f**). In contrast, introducing either D123A or M125E/Q128E into the S382A background abolished secretion (**Fig. 6e, f**). This result suggests two conclusions: first, that while this calcium site is important for secretion for both forms, it is strictly required for the secretion of proPC1/3^S382A^; and second, that the secretion of proPC1/3^S382A^ is dependent on the dimer interface. We propose that WT PC1/3 folds and matures through a unimolecular mechanism, analogous to furin and, likely, to the other PCSKs. However, when the proPC1/3 molecule is unable to mature (in this case due to the S382A mutation), it dimerizes. Dimerization represents an alternative path to complete the calcium site-I and thereby licenses ER exit (**Fig. 6g**).

## Discussion

This study establishes the chelation of the conserved calcium site-I (also known as the calcium A site) as a gatekeeping event that controls the exit of PC1/3 from the ER. By defining this chelation event as a discrete molecular step, our work bisects autocatalytic maturation and ER exit, both of which could be points for biological regulation. This study began with the unexpected observation that proPC1/3^S382A^ was able to exit the ER of HEK 293T and AtT-20 cells (**Fig. 1**). Previous reports offered conflicting results as to whether catalytically dead proPC1/3 could escape the ER of HEK 293 and AtT-20 cells.^48, 49^ In our hands, however, catalytically dead proPC1/3^S382A^ is able to efficiently exit the ER in both lines. Mechanistically, ER exit is licensed by an unanticipated domain-swapped architecture of proPC1/3^S382A^ that allows the zymogen form of proPC1/3 to achieve an exit-competent conformation. Two lines of evidence suggest that the domain-swapped dimer is not normally experienced by WT proPC1/3: 1) we did not observe maturation of proPC1/3^S382A^ *in trans* by WT proPC1/3 (**Fig. 6c**); and 2) mutations to the dimer interface prevent secretion when introduced into proPC1/3^S382A^ but not WT PC1/3 (**Fig. 6e-f**). However, when autocatalysis is blocked, as with the S382A mutation, the ability to adopt this domain-swapped dimer conformation allows proPC1/3^S382A^ to acquire the mark of a mature PC1/3 by completing its calcium-I site, bypassing the requirement for autocatalysis, and exit the ER (**Fig. 6g**).

An implication from this work is that the secretory pathway quality control machinery evaluates the structural state generated by calcium chelation following autocatalytic maturation. The components and mechanisms involved in such surveillance remain unknown. One possible scenario is that calcium chelation enables an interaction with ER export machinery. There is precedence for this scenario with PCSK9 using SURF4 and cofactors to recruit COPII machinery.^50, 51^ The ER exit of PCSK9 requires autocatalysis, but this can be overcome when the prodomain is expressed *in trans*,^34^ much like our PC1/3^ecto^ (**Supplementary Fig. 4d**). Another scenario is that calcium chelation is necessary for the protein to pass quality control surveillance for proper folding. Although the linker Asp is dispensable for autocatalytic maturation (**Fig. 6e**), whether the calcium pocket is occupied by a partially coordinated ion remains to be determined. If the pocket is unoccupied, this region of the peptidase domain is likely to be unstable. Future work will be needed to distinguish between these two scenarios, which may also inform routes to rescue the folding defects produced by disease mutations.

Whether this calcium pocket ER exit rule is generalizable to the other PCSKs remains to be tested. Notably, proPC2 exits the ER as a zymogen, protected by its dedicated chaperone, 7B2.^17, 18, 52^ Recent work has implicated ER-associated degradation in maintaining PC2 activity in α cells, highlighting the importance of ER quality control for the PCSKs.^53^ Inasmuch as the proPC1/3^S382A^ dimer is stabilized by the linker region between the peptidase and prodomain (**Fig. 3**), linker length and surface complementarity between peptidase domains likely control whether a proPC can dimerize. PC1/3 contains a longer linker than furin; however, the length of the PC1/3 linker is not an outlier within the PCSK family (**Supplementary Fig. 6a**). Previous work showed that furin’s D115-Ca^2+^ interaction is necessary to pull the cleaved P’ residues (the residues C-terminal to the primary cleavage site) out of the substrate cleft and that longer linkers in furin prevented enzymatic activity by leaving the P’ residues trapped.^54^ This suggests a trade-off between efficient ER exit and optimal enzymatic activity, highlighting the need to better understand how PCSKs are regulated. The gatekeeping nature of this particular calcium site is further supported by seminal work in which bacterial subtilisin was engineered to fold independently of its prodomain through the deletion of the loop that creates this calcium pocket.^55, 56^ Additionally, calcium site-I was the most EDTA-resistant site in furin crystals,^35^ suggesting it is highly stable once formed.

There are limitations to this study. We primarily utilized FLAG-tagged proteins expressed in and secreted from HEK 293 cells. Although we confirmed that FLAG-tagged proPC1/3^S382A^ can exit the ER in AtT-20 cells (**Fig. 1c**) and that the proPC1/3^S382A^ interaction was observed between proteins with different tags (**Fig. 3c**), it is possible that the C-terminal FLAG tag influenced the behavior of the purified proteins. Moreover, it is plausible that different folding environments (i.e., differential expression of ER chaperones) in the ER of different cells could bias the tendencies of proPC1/3 towards or against dimerization. Future work will address these questions and whether the other PCSKs are regulated by a similar mechanism.

## Resource Availability

### Lead Contact

Requests for further information and resources should be directed to the lead contact, Daniel L. Kober

### Materials Availability

All unique/stable reagents generated in this study are available from the lead contact upon reasonable request.

### Data and code availability

- The 3-dimensional cryo-EM density maps have been deposited in the Electron Microscopy Data Bank under the accession numbers EMD-77557 [https://emdataresource.org/EMD-77557]. Atomic coordinates have been deposited in the Protein Data Bank under the accession number 36GV [https://doi.org/10.2210/pdb36gv/pdb]. They are publicly available as of the date of publication.
- This paper does not report original code.
- Any additional information required to reanalyze the data reported in this paper is available from the lead contact upon reasonable request.

## Acknowledgments

This work was supported in part by the Welch Foundation (I-2246-20250403 to D.L.K.), and the UTSW Endowed Scholars Fund (D.L.K.). P.A.H. was supported by the UT Dallas Green Fellows Program. A.V.B. was supported by the UT Southwestern Lacey Graduate Fellowship.

We thank Nabil Seidah for discussions and for the kind gift of plasmids encoding mouse PC5A and PC7. We thank Chad Brautigam and Shih-Chia Tso for assistance with mass photometry experiments. Some data presented in this report were acquired with a mass photometer that was supported by award S10OD030312-01 from the National Institutes of Health. All cryo-EM data was collected at the UT Southwestern Cryo-Electron Microscopy Facility (CEMF). We thank Dan Stoddard, Ph.D., and the CEMF staff for assistance with cryo-EM data collection. The CEMF is supported by a core facilities award from the Cancer Prevention & Research Institute of Texas (CPRIT RP220582). We thank members of the Kober lab for their critical comments and suggestions on this study.

## Author Contributions

P.K. purified proteins, carried out structural studies, generated constructs, and performed cellular and enzymatic assays. P.A.H. purified proteins, performed enzymatic assays, and carried out cellular assays. R.N. performed the AtT-20 immunofluorescence experiments. A.V.B. performed enzymatic assays. C.A.P. performed cellular assays. I.L. supervised experiments, supplied reagents and contributed to writing. D.L.K. carried out structural studies, generated constructs, performed cellular assays, and wrote the manuscript with contributions from I.L. All authors edited the manuscript.

## Methods

### Plasmids and cloning

All expression constructs were generated in the pEZT vector.^57^ The parental PCSK1-3xFLAG was generated from gBlock double-stranded DNA fragment (IDT) and incorporated into the pEZT vector using Gibson Assembly. Plasmid pEZT-PCSK1-3xFLAG contains, beginning from the N-terminus, codon-optimized *PCSK1* (Uniprot P29120-1, 1-753) followed by three copies of the FLAG motif separated by two Gly-Ser linkers (DYKDDDDKGSDYKDDDDKGSDYKDDDDK, hereafter 3xFLAG). This plasmid was used as the template for site-directed mutagenesis, except for the G593R mutant, which was generated as a gBlock and assembled into the pEZT vector using Gibson Assembly methods. pEZT-PCSK1-V5, pEZT-PCSK1^S382^-V5, and pEZT-PCSK1^S382^-V5-KDEL were likewise derived from the WT or S382A mutant pEZT-PCSK-3xFLAG. The V5 tag contains the sequence GKPIPNPLLGLDST, and the V5-KDEL tag contains the sequence GKPIPNPLLGLDSTKDEL.

The pEZT-PC1/3 prodomain construct encodes human PCSK1 residues 1-110 with no epitope tag. The pEZT-PC1/3^ecto^ construct encodes, beginning from the N-terminus, human PCSK1 residues 1-27 (signal peptide) followed by residues 120-753 followed by the 3xFLAG epitope tag.

The plasmid pEZT-furin-3xFLAG contains, beginning from the N-terminus, a codon-optimized furin ectodomain (residues 1-715, Uniprot P09958), followed by the 3xFLAG epitope tag. The D153A point mutation was introduced using site-directed mutagenesis.

Plasmids encoding the open reading frames for PC7, proPC7^D187A^, and mouse PC5A were kind gifts from Dr. Nabil Seidah. These plasmids were subcloned into the pEZT and include the 3xFLAG tag at the C-terminus. The full-length mouse PC5A and residues 1-775 of PC7 or proPC7^D187A^ were amplified using PCR and assembled into a pEZT vector to encode the PCSK ectodomain followed by the 3xFLAG tag. For mouse PC5A, the S388A point mutation was introduced using site-directed mutagenesis.

The plasmid pEZT-His_10_-proSAAS contains, beginning from the N-terminus, a codon-optimized sequence for the human proSAAS signal peptide (residues 1-33), ten His residues (His_10_), followed by human proSAAS residues 34-260. The plasmid was generated using a gBlock and Gibson Assembly methods.

### Cell culture

GnTI-deficient HEK 293S cells were obtained from ATCC (CRL-3022). Cells were thawed in Dulbecco’s modified Eagle’s medium (DMEM) (high glucose) medium supplemented with 10% v/v fetal calf serum, 100 units/ml penicillin, and 100 µg/ml streptomycin sulfate and passaged in monolayer culture at 37°C in 5% CO_2_. After expanding to ten 10-cm dishes, cells were sloughed off and transferred to suspension culture in FreeStyle 293 medium (ThermoFisher) supplemented with 2% v/v FCS, 100 units/ml penicillin, and 100 µg/ml streptomycin sulfate. Suspension cells were grown with orbital shaking at 130 rpm in baffled flasks at 37°C, and 8% CO_2_. Cells were passaged to maintain a suspension density within the range of 0.4 x 10^6^ cells/ml to 3.0 x 10^6^ cells/ml.

HEK 293T cells were obtained from ATCC (ATCC CRL-3216) and were maintained in monolayer culture at 37°C and 5% CO_2_ in DMEM (high glucose) supplemented with 10% v/v FCS, 100 units/ml penicillin, and 100 mg/ml streptomycin sulfate.

AtT-20 cells were maintained in monolayer culture at 37 °C and 5% CO_2_ in DMEM (high glucose) supplemented with 10% v/v FBS.

Sf9 cells purchased from ThermoFisher (Cat # 12659017) were grown in suspension in SF900 III SFM media (Gibco) at 27°C while shaking at 130 rpm.

### Immunofluorescence

AtT-20 cells were plated on 12 mm glass coverslips at low density to achieve approximately 50% confluence at the time of fixation. 24 h after plating, cells were transfected with 0.5 µg/well pEZT-PCSK1-3xFLAG or pEZT-PCSK1^S382A^-3xFLAG plasmid using FuGENE HD transfection reagent as per the manufacturer’s instructions. AtT-20 cells transiently expressing PC1/3-3xFLAG or proPC1/3^S382A^-3xFLAG for 24 h were briefly rinsed with PBS before fixation with 4% paraformaldehyde for 15 min at room temperature, followed by 3 further rinses with PBS. Cells were permeabilized and blocked for 30 min at room temperature with 5% FBS, 0.5% Triton X-100, and 100 mM glycine in PBS. Primary antibodies were incubated overnight at 4 °C, and secondary antibodies were incubated for 1 h at room temperature. Following each antibody incubation, coverslips were washed in PBS for 5 min with gentle agitation, then mounted onto slides with Fluoromount-G (Invitrogen 4958). Primary antibodies were rabbit anti-FLAG (1:1000, Sigma-Aldrich F7425) and sheep anti-TGN38 (1:100, Serotec AHP499G). Corresponding secondary antibodies were DyLight 488 donkey anti-rabbit (1:1000, Jackson ImmunoResearch 711-485-152) and Cy3 donkey anti-sheep (1:1000, Jackson ImmunoResearch 713-165-003). Nuclei were counterstained with DAPI.

Three-dimensional image stacks were acquired on a Leica SP8 confocal microscope using a 63x oil-immersion objective at Z-step intervals optimized to the Nyquist-Shannon sampling criterion. Laser power and gain were held constant across all images. FLAG-PC1/3 and TGN38 signals were segmented and rendered as three-dimensional surfaces in Imaris 11.0.1 (Oxford Instruments) using absolute intensity thresholds that were held constant across all analyzed images.

### Protein expression and purification

BacMam baculovirus were produced in Sf9 cells as described previously.^57^ The expression of secreted proteins was induced by infecting GnTI-deficient HEK 293S cells with BacMam baculovirus using a multiplicity of infection of ∼ 3 virions per cell when the cells were at a density of 3 million cells per ml culture. The culture media was supplemented with sodium butyrate to a final concentration of 5 mM, and the cells were switched to orbital shaking at 130 rpm at 30°C and 8% CO_2_. Cells were cultured for 72 h following infection, after which cells were pelleted by centrifugation at 4°C and 4000 x g for 30 min and clarified by filtration using 0.2-micron filter units. Supernatants were stored at-80°C until purification.

To purify FLAG-tagged proteins, the conditioned supernatants were thawed and supplemented with CaCl_2_ to a final concentration of 5 mM. Conditioned supernatants were passed over anti-FLAG M2 Affinity Gel (Millipore-Sigma A2220). Resins were washed with buffer consisting of 150 mM NaCl, 50 mM HEPES-NaOH pH 6.8, and 5 mM CaCl_2_. Bound proteins were eluted in the same buffer supplemented with 400 µg per mL of FLAG peptide. Eluted proteins were further purified by gel over a Superdex 200 10/300 Increase column (Cytiva) equilibrated in buffer consisting of 150 mM NaCl, 50 mM HEPES-NaOH pH 6.8, and 5 mM CaCl_2_ operating at 4°C and a flow rate of 0.7 mL/min.

### Proglucagon cleavage assay

Proglucagon was produced in bacteria and purified by His-tag chromatography, as previously described.^28^ Recombinant proglucagon was resuspended in DMSO at 12.5 μg/μL and diluted 1:9 into the assay buffer prior to use, to make a working stock of 1.25 μg/μL. Prior to the addition of proglucagon, 100 nM enzyme was incubated in 50 mM MES pH 5.5 or 50 mM HEPES pH 7.0, 150 mM NaCl, 5 mM CaCl₂, and 0.05% Brij at 37°C for 45 min. Then, 2 μL of the proglucagon stock was added to 14 μL of the enzyme sample. The reaction mixture was incubated at 37°C for 2 hours. Reactions were terminated by adding 5x SDS loading buffer (25% (v/v) glycerol, 10% (w/v) SDS, 0.25 M Tris pH 6.8, 3% (v/v) beta-mercapto-ethanol, and 0.2% (w/v) bromophenol blue) to a 1x concentration. Samples were separated by SDS-PAGE on 15% polyacrylamide gels and visualized by Coomassie Brilliant Blue staining. Data are representative of 3 independent experiments.

### Fluorogenic protease assay

The fluorogenic pyr-RTKR-AMC substrate was purchased from VWR (Cat # I-1650.0025BA). The proSAAS CT peptide (Pen-Len) was synthesized by LSUHSC Core services. Protease activity was measured using the fluorogenic peptide substrate pyr-RTKR-AMC. Prior to activity measurements, purified PC1/3 was incubated at 100 nM in assay buffer (50 mM MES pH 5.5, 150 mM NaCl, 5 mM CaCl₂, and 0.05% Brij) at 37°C for 45 min. Following this treatment, 25 µL of enzyme samples were transferred to black, flat-bottom 96-well microplates. Reactions were initiated by the addition of 25 µL of pyr-RTKR-AMC in assay buffer to the desired final concentration for a final reaction volume of 50 μL (2%v/v DMSO final). Fluorescence was monitored continuously using a Cytation 5 imaging reader (Biotek) with excitation and emission wavelengths of 370 nm and 450 nm, respectively, using a detector gain of 65. Fluorescence measurements were collected every 60s for 30 min at 25°C. Fluorescence was converted to specific activity using a standard curve of free AMC dye. All reactions were carried out with three technical replicates. Michaelis-Menten substrate concentration parameters were obtained by calculating initial velocity (defined as the first 10 min of the reaction) using 50 nM enzyme to cleave the substrate peptide at a concentration of 6.25 to 200 µM. Michaelis-Menten data was calculated using protein from two independent purifications, with one preparation contributing three independent data points and the other contributing two.

The pH dependence curve was carried out in a similar fashion, except that the assay buffer was modified to include 150 mM NaCl, 5 mM CaCl₂, and 0.05% Brij, with either 50 mM acetate (for pH 4.5–5.0), MES (for pH 5.5–6.5), and HEPES (for pH 6.8– 7.5). 50 nM enzyme was used to cleave the fluorogenic substrate, included at a concentration of 200 µM. The pH dependence data includes four biological replicates, each the mean of three technical replicates.

### Mass photometry

Mass photometry was performed by making 100 nM stocks of the protein in a buffer containing 50mM HEPES-NaOH pH 6.8, 150mM NaCl, and 5mM CaCl_2_. 1.8 μL of the stock was added to 16.2 μL of the same buffer that had been positioned on a glass cover slip mounted and focused on a Refeyn TwoMP mass photometer. An interferometric movie of a small portion of the cover slip was taken for 60s. The movie was converted to a ratiometric format in DiscoverMP, and contrast events were related to masses using a previously obtained standard curve with BSA. The figures were rendered in DiscoverMP.

### Secretion assays

HEK 293T cells were set up on day 0 at the number and density indicated in Figure Legends. The next day, cells were transfected X-tremeGENE HP (Roche) at a ratio of 1:1.5 DNA:reagent in OptiMEM according to the instructions of the manufacturer. Transfected cells were cultured for one day, after which the conditioned media and cell fractions were processed for immunoblot. The conditioned media samples were centrifuged at 20,000 x g at 4°C for 10 min to pellet any cellular debris and the clarified supernatants were collected and subjected to SDS-PAGE. The cellular fractions were solubilized in a lysis buffer consisting of 10 mM Tris-HCl (pH 6.8), 100 mM NaCl, 1% (w/v) SDS, 1 mM EDTA, and 1 mM EGTA. Cells were lysed by 10 passages through a 23-gauge needle and subjected to SDS-PAGE followed by immunoblot analysis.

### Co-immunoprecipitation assays

HEK 293T cells were set up on day 0 in 10 cm plates at a density of 3 million cells per dish. The next day, cells were transfected with 6 µg DNA and X-tremeGENE HP (Roche) at a ratio of 1:1.5 DNA:reagent in OptiMEM according to the instructions of the manufacturer. Cells were allowed to express proteins for two more days, after which the supernatant was collected and clarified by centrifugation at 4°C and 2000 x g for 10 minutes. One percent of the supernatants was retained for immunoblot analysis and the remainder was incubated while rotating at 4°C with 20 µL equilibrated anti-FLAG Affinity Gel for 2 hours. Following this incubation, resin was pelleted by centrifugation at 4°C and 1000 x g for 2 min and washed by resuspension in 1 mL of buffer consisting of 150 mM NaCl, 50 mM HEPES-NaOH pH 6.8, and 5 mM CaCl_2_, followed by rotation at 4°C for 30 min. The resin was washed three times before bound proteins were eluted in 400 µL of 200 mM glycine pH 3.5. The retained input and collected eluates were subjected to SDS-PAGE and immunoblot analysis.

### Immunoblot

All immunoblot experiments are representative of at least three independent experiments with similar results. Following SDS-PAGE, proteins were transferred to nitrocellulose membranes using the Trans-Blot Turbo Transfer System (Bio-Rad Laboratories, Hercules, CA). Membranes were probed with the primary antibodies (anti-FLAG, Millipore-Sigma Cat #F1804, 1:1000; anti-V5, Sigma Cat # V8012, 1:5000; or anti-Calnexin, Fisher Cat # 50-172-6300, 1:2000). Bound antibodies were detected by chemiluminescence (SuperSignal West Pico Chemiluminescent Substrate, Thermo Scientific, Waltham, MA) using a 1:5000 dilution of anti-mouse or a 1:10,000 dilution of anti-rabbit IgG conjugated to horseradish peroxidase (Jackson ImmunoResearch Laboratories, Inc., West Grove, PA). Membranes were exposed to Blue X-ray Film (Phoenix Research Products, Pleasanton CA).

### Sequence alignments

The following human PCSK sequences were aligned: PCSK1 (Uniprot P29120), PCSK2 (Uniprot P16519), FURIN (Uniprot P09958), PCSK4 (Uniprot Q6UW60), PCSK5 (Uniprot Q92824), PCSK6 (Uniprot P29122), PCSK7, (Uniprot Q16549), MBTPS1 (Uniprot Q14703), and PCSK9 (Uniprot Q8NBP7). Clustal Omega was used to produce the alignments and phylogenetic data.^58^ MEGA 12^58^ was used to generate the phylogenetic tree using outputs from Clustal Omega. The ESPript 3.2 server^59^ was used to visualize the aligned sequences of the kexin-like PCSKs using outputs from Clustal.

### Cryo-EM

#### Cryo-EM grid prep

Quantifoil R 1.2/1.3 300 Mesh, Au grids were glow discharged for 80 seconds at 30 mA using a Pelco easiGlow machine. 4 µL sample at 0.4 mg/mL was applied at 4°C in 100% humidity and blotted for 4.5 s followed by freezing in liquid ethane cooled by liquid N2, using a Vitrobot Mark IV.

#### Cryo-EM Data Collection

6285 Movies were collected using a Titan Krios G2 operating at 300 kV using a K3 Summit direct electron detector (Gatan) in the super-resolution counting mode using SerialEM software.^60^ The super-resolution pixel size was 0.413Å, the dose rate was ∼ 17 e-/Å^2^/sec and the total accumulated dose was 50 e-/ Å^2^ over 50 frames. Data were acquired on a 3×3 pattern of holes with three movies collected per hole using image shift. A nominal defocus range of-0.8 to-2.0 microns was used, and the energy filter was set to 20 eV.

#### Cryo-EM image processing

Initial processing was carried out using CryoSparc v4.7.0.^61^ Movies were subjected to Patch Motion Correction, where they were Fourier cropped to a pixel size of 0.826 Å, followed by Patch CTF Estimation. A curated subset of 6139 micrographs was selected using a criterion of CTF fit to 4.5Å. Gaussian blob picking was performed on 100 micrographs to generate quality reference images for template picking. Six million particles resulting from template picking were extracted in a box size of 300 pixels that were Fourier cropped to 128 pixels. These particles were subjected to one round of 2D classification to remove obvious junk particles, resulting in 1,768,115 particles. A random subset of 600,000 particles was used to generate four *ab-initio* models, which were used to sort the full dataset using Heterogeneous 3D refinement. A single good class of 765,175 particles were re-extracted in a box of 300 pixels and cropped to 256 pixels. Non-uniform 3D refinement (NU3D) yielded a map with a nominal resolution of 2.95Å that suffered obvious defects from anisotropic resolution. 3D classification without alignment yielded a best class with 145,682 particles that produced a higher-quality map with nominal resolution of 3.0 Å by NU3D. Application of C2 symmetry improved the map to 2.9 Å. Finally, 143,577 particles were re-extracted in boxes of 340 pixels that were cropped to 300 pixels, resulting in a pixel size of 0.936 Å. NU3D of these particles resulted in a map of 2.65Å, however, this map and a subsequent 2.48 Å map generated following Reference-Based Motion Correction proved difficult to model due to noisy features. Instead, the particles were subjected to 3D auto-refinement in Relion v5 using Blush Regularization.^62^ This refinement resulted in a map with a nominal resolution of 2.9 Å that could be confidently modeled. Relion v5 was obtained through the SBGrid Consortium.^63^ For model building, the map was filtered by local resolution using a B factor of-50. Figure Legends indicate whether the sharpened map or unsharpened map are used for each illustration.

#### Model building

Initial models were obtained using AlphaFold 3 to predict the proPC1/3 dimer interaction. Starting models were subjected to rounds of manual building in Coot^64^ and ISOLDE^65^ followed by real-space refinement in Phenix.^66^ The final refinements incorporated PDB 9FIC (high resolution X-ray structure of the PC1/3 prodomain) as a reference model. Refinement statistics are provided in **Supplemental Table 1**. The atomic model includes residues 30-602, except for one flexible loop that was omitted (residues 134-143).

## Statistical Analysis

Immunoblot densitometry was performed using ImageJ. One-way ANOVA analysis was carried out using GraphPad Prism 10 using Tukey’s method to correct for multiple comparisons. Enzymatic analysis was carried out using GraphPad Prism 10.

