## supplemental figures and table for "A domain-swapped proPC1/3 structure reveals a conformational checkpoint for ER export"

**Supplemental Table 1. Cryo-EM data collection, refinement and validation statistics**

| | proPC1/3$^{S382A}$ |
|---|---|
| **Data collection and processing** | |
| Magnification | 105,000 |
| Voltage (kV) | 300 |
| Electron exposure (e–/Å$^2$) | 50 |
| Defocus range (μm) | -0.8 to -2.0 |
| Pixel size (Å) | 0.826 |
| Symmetry imposed | C2 |
| Initial particle images (no.) | 1,768,115 |
| Final particle images (no.) | 143,577 |
| Map resolution (Å) | 2.9 |
| FSC threshold | 0.143 |
| Map resolution range (Å) | 2.9 - 4.2 |
| **Refinement** | |
| Initial model used (PDB code) | AlphaFold 3 |
| Model resolution (Å) | 3.1 |
| FSC threshold | 0.5 |
| Model resolution range (Å) | - |
| Map sharpening *B* factor (Å$^2$) | -50 |
| Model composition | |
| Non-hydrogen atoms | 8,878 |
| Protein residues | 1,126 |
| Ligands | NAG: 4 |
| | CA: 6 |
| | NA: 2 |
| *B* factors (Å$^2$) | |
| Protein | 71.70 |
| Ligand | 72.86 |
| R.m.s. deviations | |
| Bond lengths (Å) | 0.005 |
| Bond angles (°) | 0.539 |
| Validation | |
| MolProbity score | 1.40 |
| Clashscore | 3.24 |
| Poor rotamers (%) | 0.21 |
| Ramachandran plot | |
| Favored (%) | 95.89 |
| Allowed (%) | 4.11 |
| Disallowed (%) | 0.00 |

### Supplemental Figures and Legends

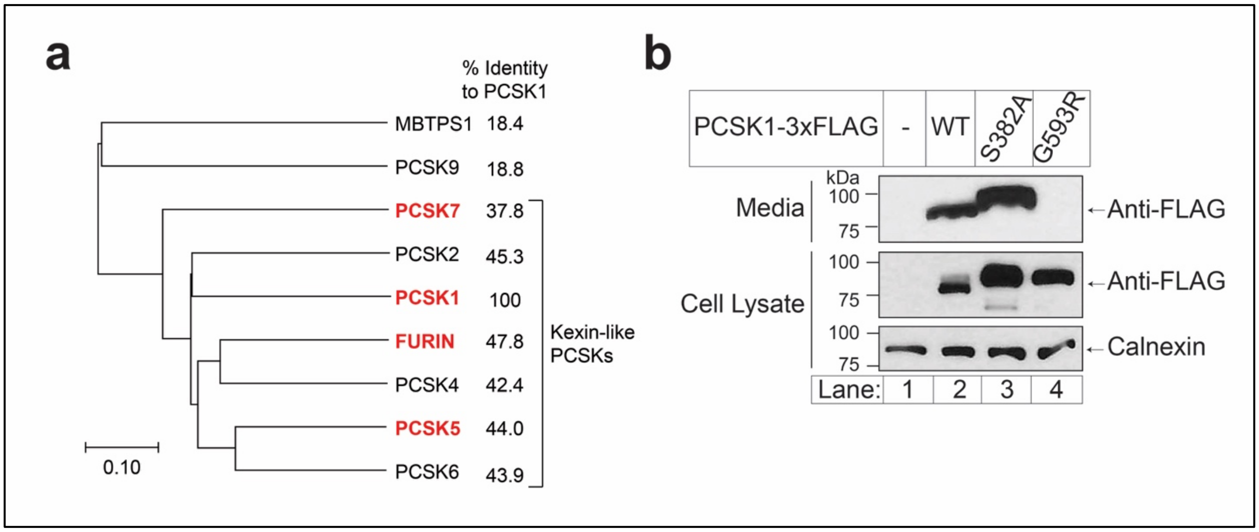

**Supplemental Figure 1. PCSK sequence identity and expression of PC1/3 variants.**

**a)** Phylogram of human PCSKs, generated using Clustal Omega and visualized using MEGA 12 software. The % identity of the amino acid sequence to PCSK1 is indicated. Bracket indicates the fractional substitutions per position. Red gene names indicate proteins tested in Figure 1, except that Figure 1 uses mouse PC5A instead of human PC5A.

**b)** HEK 293T cells do not secrete the PC1/3 disease variant G593R. HEK 293T cells were set up in 12-well dishes at a density of 400,000 per well. The next day, cells were transfected with 1 µg of empty pEZT vector (lane 1) or the indicated plasmid (lanes 2-4). Samples were electrophoresed on 10% SDS-PAGE and subjected to immunoblot analysis for the indicated epitopes as described in Methods.

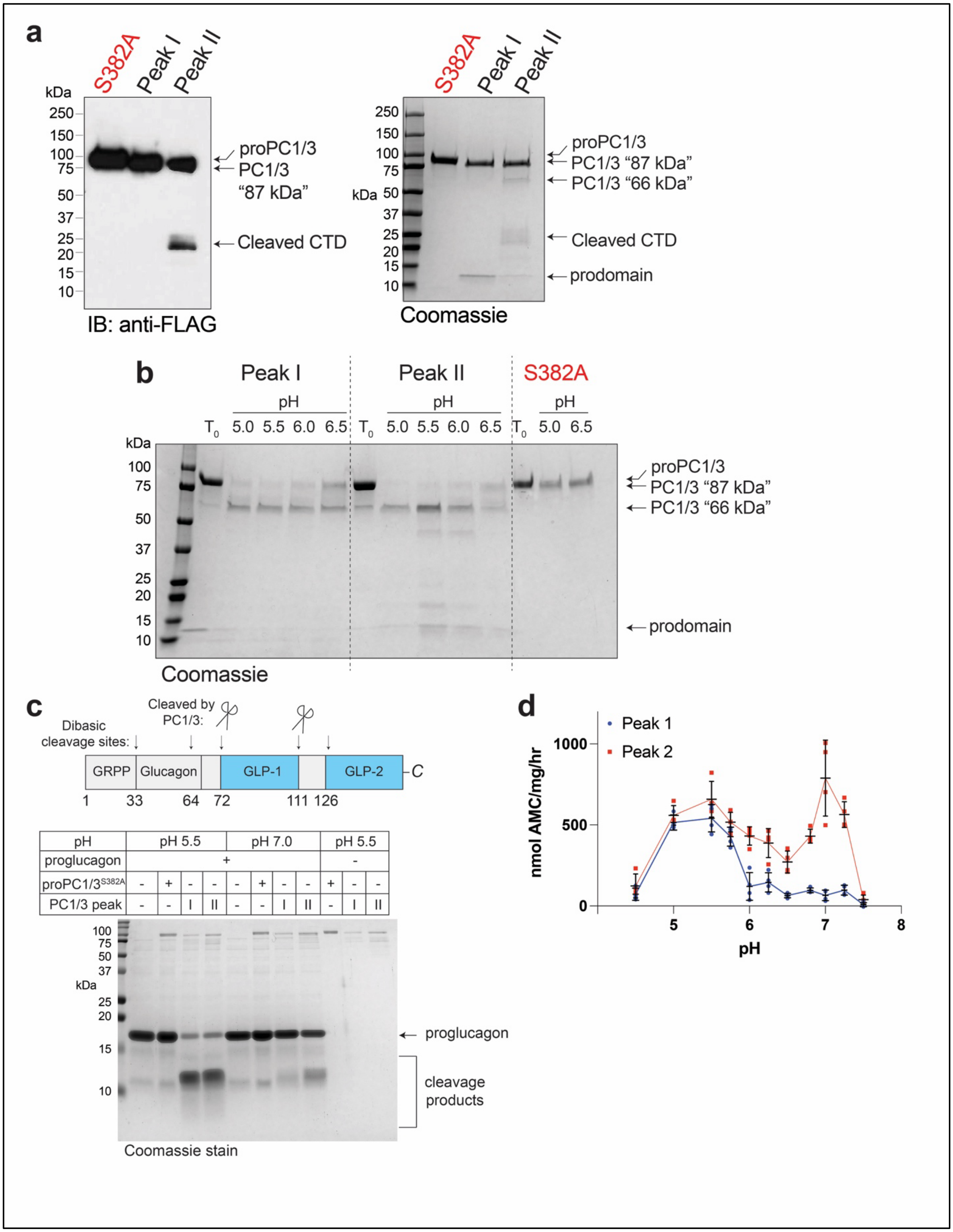

**Supplemental Figure 2. Biochemical and enzymatic characterization of purified human PC1/3 and proPC1/3$^{S382A}$.**

**a)** Analysis of purified samples by anti-FLAG immunoblot (left) and Coomassie-stained SDS-PAGE (right) for purity and to identify cleavage products. Both samples were electrophoresed on 4-15% SDS-PAGE.

**b)** Catalytic maturation of full-length PC1/3 into its 66 kDa form. Purified PC1/3 and proPC1/3$^{S382A}$ (see **Fig. 2b**) were incubated overnight at room temperature in protease assay buffer adjusted to the indicated pH. Samples were electrophoresed on 4-15% SDS-PAGE and visualized by Coomassie-staining. Representative of three independent experiments.

**c)** Cleavage of proglucagon by purified PC1/3 Peak I and Peak II. 2.5 µg of recombinant His-tagged proglucagon was processed by PC1/3 at the indicated pH as described in Methods. Reaction products were separated by 15% SDS-PAGE and visualized by Coomassie stain. Representative of three independent experiments.

**d)** The pH dependence of the cleavage of pyr-RTKR-AMC by PC1/3 Peak I and Peak II. Buffers used to adjust pH were sodium acetate (pH 4.5 and 5.0), MES (pH 5.5, 5.75, 6.0, 6.25, and 6.5), and HEPES (pH 6.8, pH 7.0, pH 7.25, and pH 7.5). Mean values (black bars) were derived from the means four independent experiments (red and blue shapes), each conducted three technical replicates. Error bars are SD.

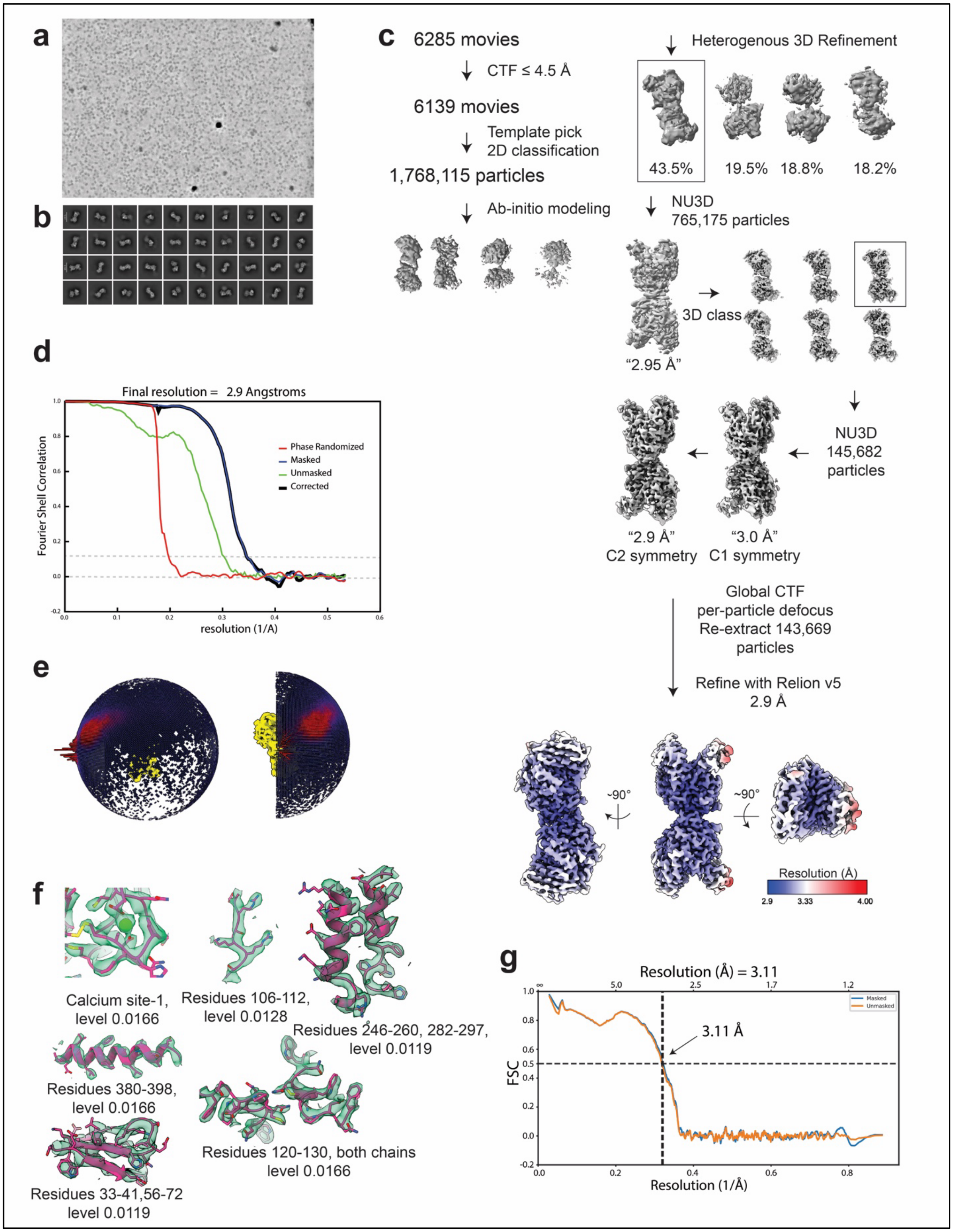

**Supplemental Figure 3. Cryo-EM data processing of proPC1/3$^{S382A}$.**
**a)** Representative denoised micrograph.

**b)** 2D classification results.
**c)** processing flowchart. All steps except the final refinement were carried out using CryoSparc v4.7. Final map is shown colored by local resolution.
**d)** GSFSC curves (2.9 Å resolution with 0.143 cut-off).
**e)** 3D viewing orientations projected onto the final map.
**f)** Representative densities from the sharpened map and structure model. Residues and map contours for each section are indicated.
**g)** Map/Model FSC (3.1 Å with 0.5 cut-off).

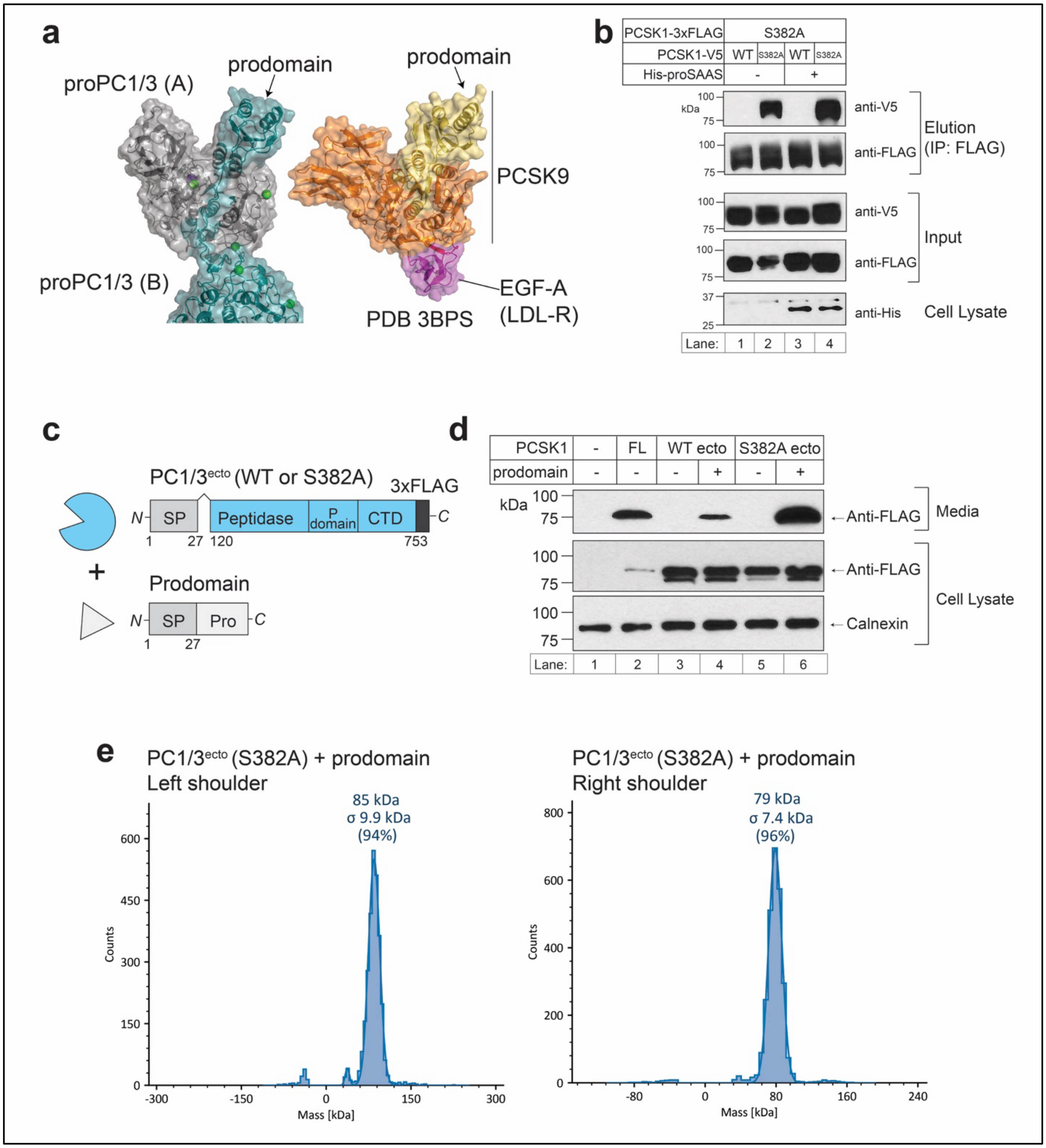

**Supplemental Figure 4: Analysis of proPC1/3$^{S382A}$ dimer interactions**

**a)** The EGF-A domain of the LDL receptor (LDLR, shown in pink) binds to PCSK9 (shown in yellow and orange) through a surface analogous to the proPC1/3$^{S382A}$ dimer interface. PDB 3BPS was aligned with the structure of the proPC1/3$^{S382A}$ dimer.

**b)** Co-expression of proSAAS does not affect the proPC1/3$^{S382A}$-proPC1/3$^{S382A}$ interaction. HEK 293T cells were set up and transfected with 2 µg each of the indicated plasmids. Empty pEZT vector was used to adjust the total DNA amount to 6 µg. Following expression for two days, samples were subjected to anti-FLAG co-IP, as described in Methods. Samples were electrophoresed on 8% SDS-PAGE.

**c)** Schematic for the PC1/3$^{ecto}$ and prodomain plasmids. Both contain the native PCSK1 signal peptide for insertion into the ER.

**d)** Rescue of PC1/3$^{ecto}$ secretion by co-expression with an untagged prodomain. HEK 293T cells were set up in 12-well plates at a density of 400,000 cells per well and transfected 0.5 µg of the indicated PC1/3$^{ecto}$ construct and 1 µg of the prodomain construct, as indicated. Empty pEZT vector was used to adjust the total amount of plasmid DNA to 1.5 µg. After expressing for one day, samples were harvested and processed for immunoblot as described in Methods.

**e)** Mass photometry analysis of the purified PC1/3$^{ecto}$/prodomain complex (samples from Fig. 3E).

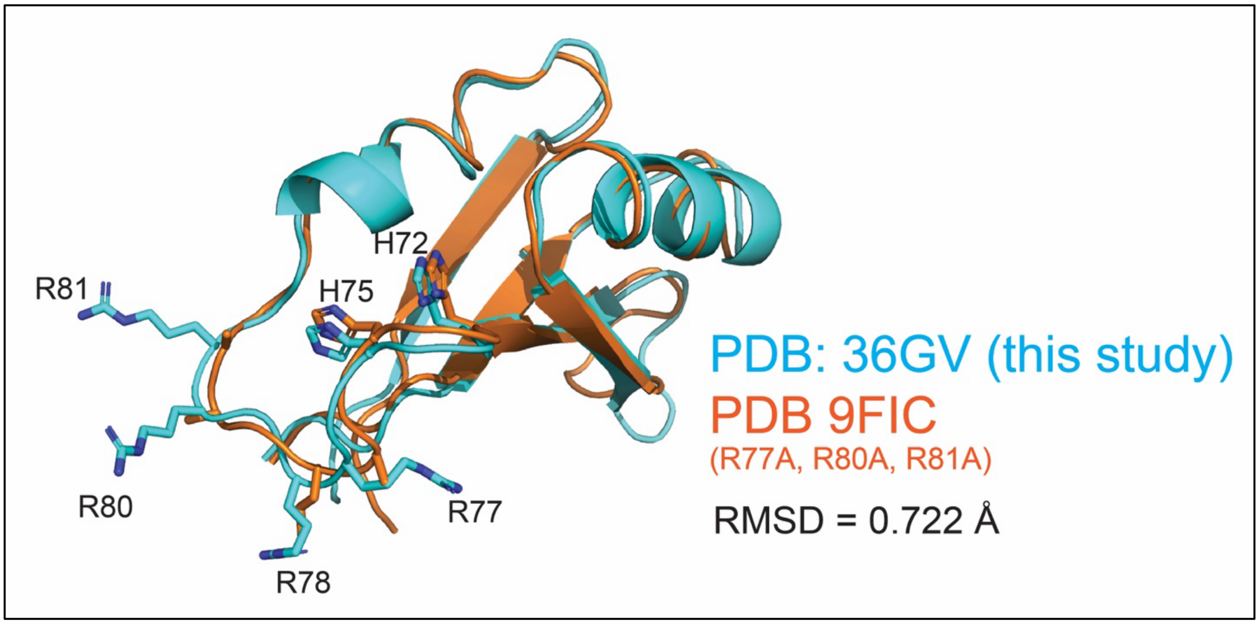

**Supplemental Figure 5: PC1/3 prodomain**

**a)** Superposed PC1/3 prodomains. Key His residues and secondary cleavage loop residues are highlighted with side chains in sticks. Panel generated with Pymol.

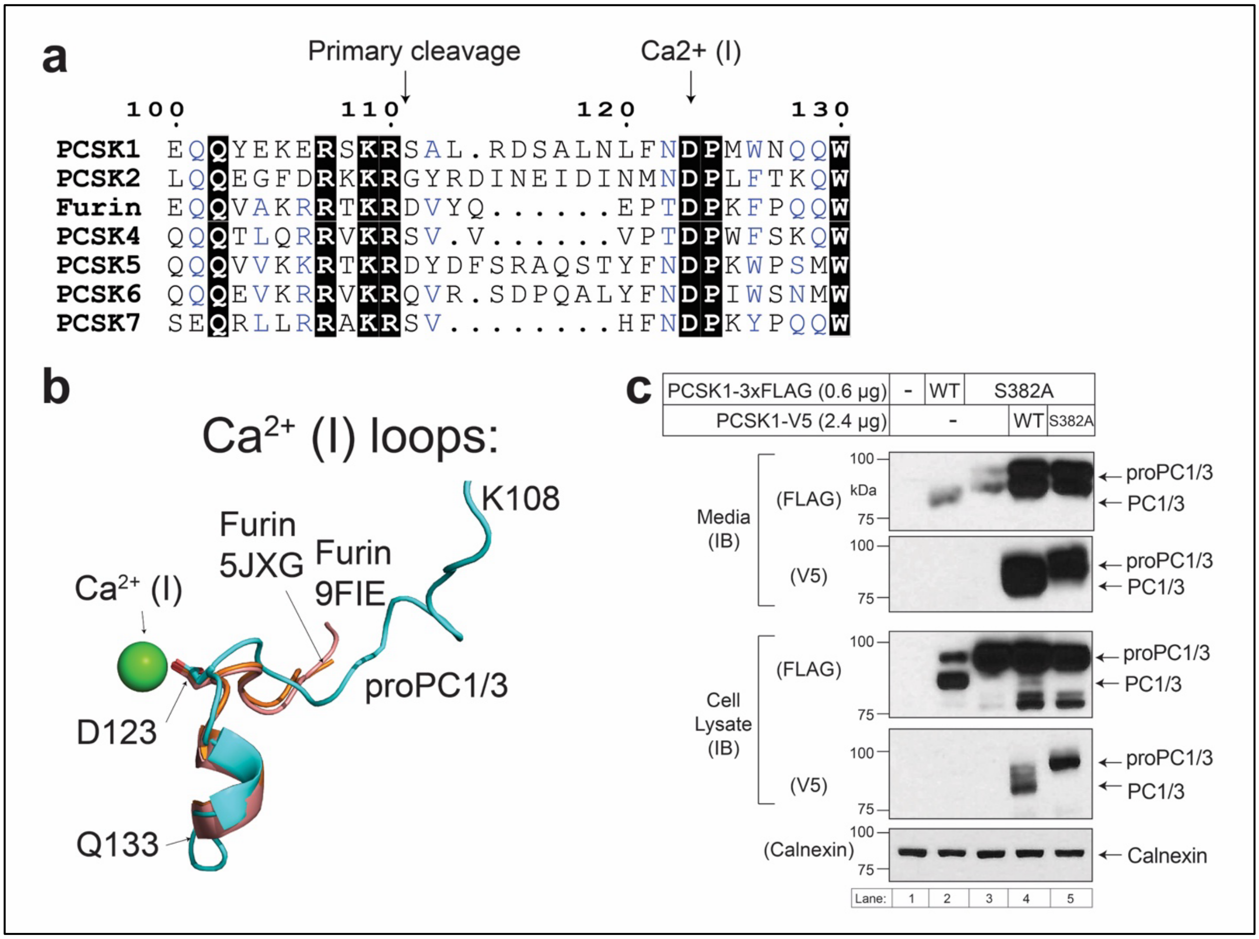

**Supplemental Figure 6: Sequence and expression analysis of proPC1/3**

**a)** Alignment of human kexin-like PCSKs. Conservation analysis visualized using ESPript server. Black boxes indicate invariant residues. Blue text highlights conserved consensus residues.

**b)** Superposed linker motifs from indicated structures. Labels indicate the coordinating Asp and the N- and C-terminal residues from this motif in proPC1/3$^{S382A}$. Two examples from mature furin are shown in brown and orange (PDB 5JXG and 9FIE). ProPC1/3$^{S382A}$ is shown in cyan.

**c)** ProPC1/3$^{S382A}$ not efficiently cleaved *in trans* by PC1/3. HEK 293T cells were set up in 6-well plates at a density of 1 million cells per well and then transfected with 3 µg of DNA as indicated or with empty pEZT vector. After expressing for one day, cell and supernatant samples were harvested and processed as described in Methods. Samples were electrophoresed on 8% SDS-PAGE. IB = immunoblot for the epitope tags indicated in parentheses.
